# Multiscale Free-Energy Methods for Protonation-Coupled Light-Responsive Binding of Ionizable Photoswitchable eDHFR Inhibitors

**DOI:** 10.64898/2026.05.16.725670

**Authors:** Mohammad Khavani, Kambham Devendra Reddy, Pauf Neupane, Gustavo J. Costa, Laleh Khalvati, Ruibin Liang

## Abstract

Photoswitchable ligands enable photocontrol of biomolecular activity by binding to targets in an isomer-dependent, light-responsive manner. Recent developments in ionizable photoswitchable ligands greatly expand their applications but introduce a major design challenge: light-responsive binding can depend on isomeric form, chemical substitution, and binding-induced shifts in protonation equilibria. These effects are tightly coupled, subtle in magnitude, and difficult to predict. Consequently, few computational methods have been developed and systematically benchmarked for quantitatively predicting them. Here, we establish a multiscale free-energy method and benchmark it against experimental data for a series of recently developed photoswitchable inhibitors of *Escherichia coli* dihydrofolate reductase (eDHFR), a crucial target in photopharmacology. Constant pH replica-exchange molecular dynamics and quantum mechanics/molecular mechanics umbrella sampling quantitatively characterize the ligand’s protonation-state change upon binding to the eDHFR active site. Thermodynamic integration simulations using alternative alchemical pathways, thermodynamic cycles, and protonation-state assignments were evaluated for predicting light-responsive affinity differentials and substituent effects. Direct *cis*-to-*trans* transformations with explicit treatment of environment-dependent protonation states best reproduce experimental trends. Compound-to-compound pathways are less reliable because force-field inaccuracies introduce large pKₐ errors that are difficult to correct when protonation/deprotonation processes implicitly enter the thermodynamic cycle. TI simulations that ignore binding-induced protonation-state changes fail to consistently reproduce experimental trends. Protein-ligand and ligand-water interaction analyses further reveal the energetic and structural origins of isomer-dependent binding. This study establishes a systematic free-energy method for designing ionizable photoswitches in photopharmacology.

## Introduction

Molecular photoswitches provide a powerful toolset for the reversible control of biomolecular systems with light.^1–4^ By switching between distinct isomeric forms, they enable noninvasive, remote, and selective photocontrol of target proteins with high spatial and temporal precision, thereby reducing off-target effects that remain a major challenge for conventional therapeutics.^5–7^ These advantages have driven the rapid expansion of photopharmacology, in which molecular photoswitches are designed as light-regulated ligands that target specific biomolecules to minimize side effects.^1–15^ The key to their successful design is achieving pronounced bioactivity contrast between the light-activated and dark-adapted states. To this end, first, the isomeric form with the highest bioactivity/affinity towards the target (referred to as “active isomer” below) needs to be correctly identified. For example, a “*cis*-on” effect indicates that the *cis* isomer of the ligand has higher bioactivity/affinity than the *trans* isomer, whereas a “*trans*-on” effect indicates the opposite. Second, the difference in bioactivity and/or binding affinity between distinct isomeric forms of the ligand needs to be maximized.

Therefore, the rational design of photoswitchable ligands can be greatly facilitated by accurate computational predictions of three quantities: the active isomer for a given biomolecular target, the light-responsive binding affinity differential between the active and inactive isomers, and the effect of chemical substitutions on this differential. Experimentally optimizing these properties often requires extensive trial and error, whereas all-atom simulations can, in principle, reveal the protein-ligand interactions that underlie isomer-specific binding. However, quantitative prediction remains challenging. The structural differences between isomers are often localized near the isomerizing bond, and the resulting binding free-energy differences are typically small, often below a few kilocalories per mole.^16^ Errors of similar magnitude can therefore lead to incorrect prediction of the active isomer and misguide ligand design.^17^ Standard molecular docking, unbiased molecular dynamics, and end-point free-energy methods often lack the accuracy required to resolve these subtle light-responsive affinity differentials and the effects of substituents on them, necessitating rigorous free-energy approaches.^18, 19^

An additional design challenge arises for ionizable photoswitchable ligands containing titratable groups, which have recently become increasingly important in photopharmacology because their photochemical and biochemical properties are tunable by pH in addition to illumination. An example is photoswitchable acylhydrazones that can tautomerize between multiple protonation states.^20^ Their dominant protonation state may differ between the bulk solution and the protein’s binding pocket, where local electrostatic environments can strongly shift protonation equilibria.^21,22^ In such cases, light-responsive binding depends not only on the geometric difference between isomers but also on how ligand-protein interactions shift protonation equilibria. Thus, accurate prediction of light-responsive binding for ionizable photoswitches requires free-energy methods that consistently treat the coupled effects of isomerization, chemical substitution, and protonation-state changes on protein-ligand interactions.

Here, we address these challenges using recently developed photoswitchable inhibitors of *Escherichia coli* dihydrofolate reductase (eDHFR) as a model system. DHFR catalyzes the NADPH-dependent conversion of dihydrofolate to tetrahydrofolate, a key cofactor required for nucleotide and amino acid biosynthesis,^23, 24^ and has long served as a platform for structure-based inhibitor design. Well-established DHFR inhibitors include methotrexate, which is widely used in cancer chemotherapy to target human DHFR^23^, as well as trimethoprim (TMP), a clinically important antibacterial agent commonly prescribed for urinary tract infections.^25, 26^ Recent work has demonstrated that DHFR activity can be effectively modulated using light^14, 17, 27, 28^ via TMP-derived photoswitchable inhibitors, in which the drug scaffold is conjugated to an azobenzene moiety. In a previous experimental design campaign^17^, a series of such compounds, including compounds **6, 11**, and **15**, was synthesized and characterized (**Figure 1A**). Compound **11** introduces a para-carboxyl substituent relative to compound **6**, whereas compound **15** contains both a meta-chloro and para-carboxyl substitution (**Figure 1B**). These modifications substantially alter both absolute inhibitory potency and the light-responsive bioactivity differential. In particular, the introduction of the carboxyl group (**6**→**11**) increases absolute affinity but unexpectedly reduces the “*cis*-on” effect, whereas further adding the chlorine substituent (**11**→**15)** partially restores light-responsive activity.

**Figure 1.**
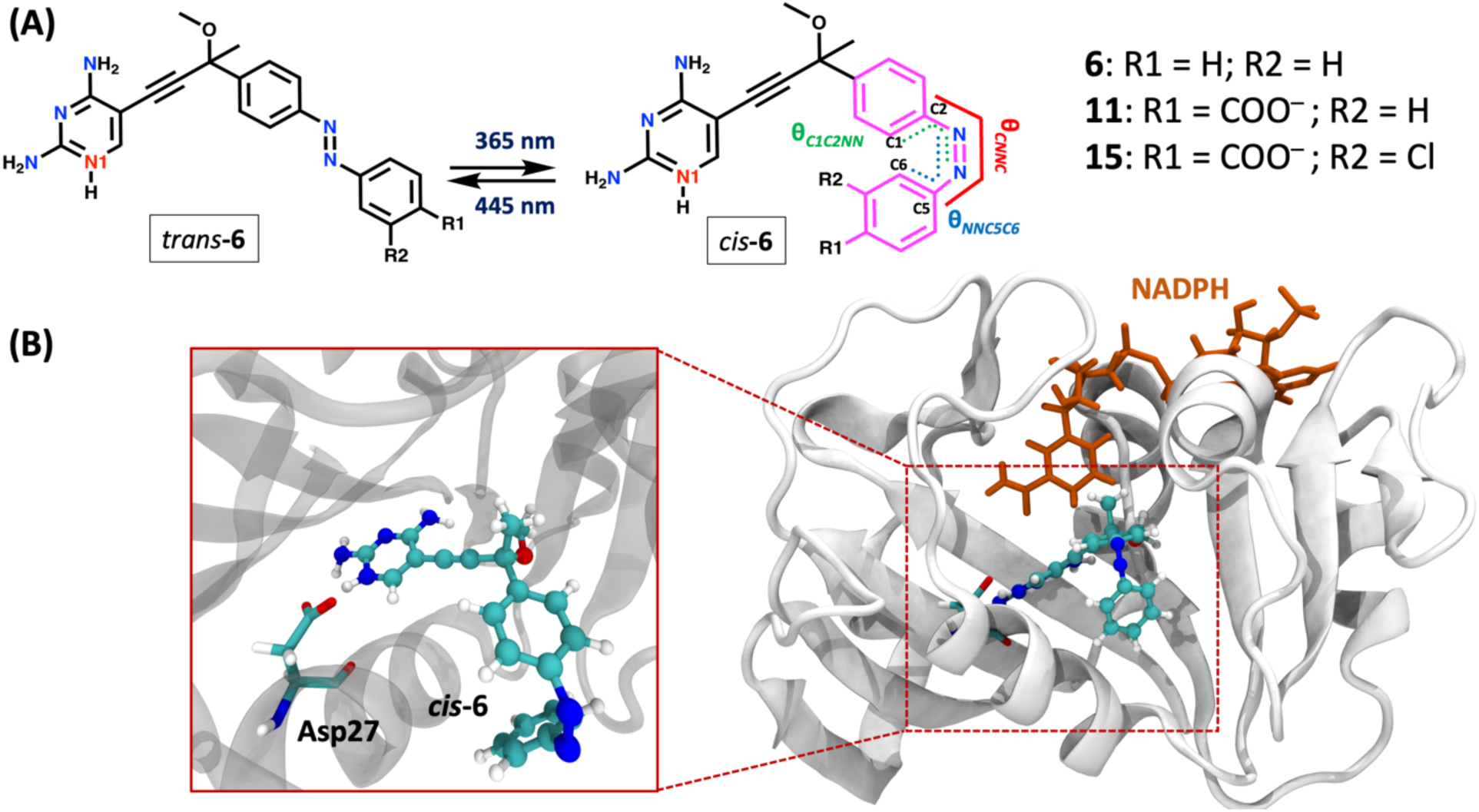
(A) Schematic illustration of the photoisomerization of TMP-derivatives (compounds **6, 11** and **15**) under illumination at different wavelengths, along with their chemical structures. The N1 atom in the diaminopyrimidine ring (in red) can be either deprotonated or protonated. (B) The eDHFR (gray ribbons) in complex with the *cis*-**6** (sticks and balls), highlighting the Asp27 residue (sticks) in the binding pocket and binding-induced protonation of the N1 atom in the diaminopyrimidine ring. The NADPH (sticks) is shown in orange.

Despite these qualitative insights from the experiment^17^, it remains challenging to computationally quantify the role of substituents’ effects on the light-responsive affinity differential of TMP-based photoswitchable inhibitors. A major challenge is that their ligand-binding affinity may be influenced by changes in protonation state upon binding to eDHFR.^29^ In particular, the pK_a_ of the diaminopyrimidine ring (**Figure 1A**), which is shared across all TMP-based photoswitchable inhibitors, can be significantly elevated through interactions with ionizable residues such as Asp27 (**Figure 1B**).^17, 29, 30^ As a result, ligand binding can induce the protonation of the N1 atom on the diaminopyrimidine ring (**Figure 1A**). The shift in this protonation equilibrium could qualitatively change the active isomer form of the photoswitchable ligand when the *cis*-vs-*trans* affinity differential is small. When rationalizing the molecular basis of *cis*-vs-*trans* affinity differential via conventional MD simulations, the previous study^17^ did not consider the potential change in protonation state upon shifting from aqueous solution to the active site. Additionally, there were no rigorous free energy calculations for the *cis*-vs-*trans* affinity differential or the substituents’ effects on this quantity. As a result, these MD simulations^17^ have not consistently predicted the effects of substituents on differential light-responsive activity. We reason that these critical missing components, if implemented in a quantitative, multiscale free-energy framework, would have benefited quantitative prediction in the previous experimental design. ^17^

Here, we establish a multiscale free-energy method to quantify protonation-coupled, light-responsive binding of these TMP-derivatives in ionizable, photoswitchable eDHFR inhibitors. Constant-pH replica-exchange molecular dynamics (pH-REMD) and quantum mechanics/molecular mechanics (QM/MM) umbrella sampling were employed to characterize ligand protonation in solution and in the eDHFR active site. These calculations define the protonation-state assignments used in thermodynamic integration (TI) simulations^31–33^, which were then benchmarked against experimental data for the *cis*-vs-*trans* affinity differential and substituent effects. We compare alternative alchemical transformation pathways, including *cis*-to-*trans* transformations for each ligand and compound-to-compound transformations within fixed isomeric forms, and evaluate the sensitivity of these strategies to different treatments of protonation states. The predictive accuracy of endpoint methods, such as Molecular Mechanics Poisson-Boltzmann/Generalized-Born Surface Area (MM-PB(GB)SA) ^34–37^ was also benchmarked. Additionally, we analyzed protein-ligand and ligand-water interactions to interpret the energetic and structural origins of isomer-dependent binding. Overall, this study establishes a systematic computational framework for designing ionizable photoswitches in photopharmacology.

## Methods

### System setup

The crystal structure of eDHFR (PDB ID: 3DAU)^38^ was selected as the starting model for all molecular docking and molecular dynamics (MD) simulations. For each TMP-based ligand in either the *cis* or *trans* isomer, AutoDock Vina^39^ was employed to place it in the substrate-binding pocket of eDHFR. The resulting binding poses were inspected to ensure that key interactions with binding-site residues were, whenever possible, consistent with those previously reported in ref ^17^. The NADPH molecule in the crystal structure was retained, while all crystal water molecules were removed because none were buried deep within the substrate-binding pocket. The initial protonation states of ionizable amino acid residues at physiological pH were assigned using the H^++^ server.^40^ Both protonation states of the ligand were set up and tested. The protein-ligand complexes were then solvated in a box of explicit water molecules under periodic boundary conditions (PBC) of ∼85 × 84 × 90 Å^3^. Water molecules were modeled using the TIP3P model^41^, while the protein was described using the AMBER ff14SB force field^42^. The parameters in ref ^43^ were used for NADPH. Ligand parameters were first generated using the general AMBER Force Field (GAFF) approach.^44, 45^ For each isomer of each ligand, a separate aqueous solution set up where the ligand was solvated in a PBC box of water molecules of ∼ 45 × 45× 45 Å^3^.

The torsional parameters in the force field for three dihedral angles, C2-N3=N4-C5 (θ_CNNC_), C1-C2-N3=N4 (θ_C1C2NN_), and N3=N4-C5-C6 (θ_NNC5C6_) (**Figure 1A**), were reparametrized using relaxed scans of their corresponding potential energy surface (PES) with a multireference electronic structure method. This step was necessary because our previous studies^18, 19, 46, 47^ indicated that unmodified GAFF parameters have artificially low torsional barriers, leading to unphysically fast *trans* ↔ *cis* thermal isomerization on nanosecond timescales during MD simulations. Due to the multireference character of the electronic wavefunction near the transition state of the θ_CNNC_ torsion,^48^ all three torsional terms were refined based on relaxed PES scans of the N1-deprotonated ligands (**Figure S1**) using the floating occupation molecular orbital hole−hole Tamm-Dancoff approximated density functional theory (FOMO-hh-TDA-DFT) electronic structure method,^49, 50^ with the BH&HLYP functional and the 6-31+G(d) basis set. This multireference electronic structure method has the advantage of incorporating both static and dynamic electron correlation and has been extensively validated for both the ground and excited-state PES and dynamics of azobenzene-based photoswitches.^19, 46–48, 50–54^ The torsional parameters were adjusted until the relaxed-scanned PES using the force field closely matched the relaxed-scanned PES using the FOMO-hh-TDA-DFT method in terms of *cis*-to-*trans* energy differences and isomerization barriers (**Figure S1**). All QM calculations used for force field refinement were performed with the TeraChem software package.^55, 56^

### Force-field-based MD equilibration simulations

Each system was initially relaxed through 100,000 steps of energy minimization, followed by gradual equilibration in the constant NVT ensemble at 300 K for 2 ns using a 1 fs time step. During this equilibration phase, positional restraints were applied to all non-hydrogen atoms of the protein using harmonic potentials with a force constant of 50.0 kcal/mol/Å^2^. Subsequently, restrained MD simulations were carried out in the constant NPT ensemble at 300 K and 1 atm, with a 1 fs time step, gradually reducing the positional restraint to 0.1 kcal/mol/Å^2^ over 5 ns. Then, removing any positional restraints, constant NPT simulations were carried out for a total of 200 ns. The initial 50 ns of each trajectory were discarded as equilibration stage, and the remaining 150 ns were used for interaction energy analysis. The Langevin thermostat^57^ with a collision frequency of 1 ps^-1^ was applied to maintain the temperature at 300 K, while the pressure was regulated by a Monte Carlo barostat.^58^ Electrostatic interactions were evaluated using the Particle Mesh Ewald (PME) method,^59^ and a cutoff distance of 12 Å was employed for van der Waals interactions. All bonds involving hydrogen atoms were constrained using the SHAKE algorithm,^60^ allowing the use of a 2 fs timestep throughout the simulations. All minimizations and MD simulations were performed with the Amber24 software package.^61^

### pH-REMD simulations

To identify the protonation state of the photoswitchable TMP derivatives in both the aqueous solution and eDHFR, pH-REMD simulations^62^ were carried out for compound **6** as a representative example. To determine the deprotonation free energy (ΔG_deprotonation_) of **6** in aqueous solution, a thermodynamic integration (TI)^32^ simulation was first performed to alchemically transform protonated **6** to deprotonated **6**. Details for this TI simulation are described in the TI section below. Then, the reference pKa value in the aqueous solution was calculated using DFT with the M06-2X functional^63^ and TZVP basis set in a conductor-like polarizable continuum model (M06-2X/TZVP/CPCM, 𝜀 = 80.15)^64^, following similar computational protocols in previous studies.^65, 66^ The reference ΔG_deprotonation_ and pKa values were then used as input parameters for the pH-REMD simulations in the aqueous solution and protein.

Prior to performing pH-REMD simulations, a multi-step equilibration protocol was applied to both the aqueous solution and protein-ligand complex systems. Each system was first subjected to energy minimization to remove unfavorable contacts. Minimization was carried out using a combination of steepest descent and conjugate gradient methods for a total of maximum 50,000 steps, with the first 10,000 steps performed using steepest descent. During this stage, harmonic positional restraints (10 kcal/mol/Å²) were applied to the ligand while allowing the environment to relax. A cutoff of 12 Å was used for nonbonded interactions, and the systems were treated under PBC with explicit solvent. Following minimization, the systems were gradually heated to 300 K using the constant NVT ensemble for 1 ns. Afterwards, a 1 ns equilibration simulation was performed in the constant NPT ensemble at 300 K and 1 atm. Positional restraints (5 kcal/mol/Å^2^) were applied to the ligand during this step. A second equilibration stage was then carried out using constant pH MD (CpHMD) at pH=4.5 without positional restraints at 300 K. In this step, the system was simulated for 1 ns for the ligand in water and 6 ns for the ligand-protein complex. Importantly, in the protein-ligand complex systems, both the Asp27 and the ligand wer treated as ionizable in the CpHMD simulations.

Following the pre-equilibration steps, pH-REMD simulations were carried out over a pH range of 1.0-7.5 for the aqueous solution (14 replicas) and 7.0-17.5 for the protein-ligand complex (22 replicas). The pH values of the replicas were separated 0.5 pH units apart, and replica exchange was attempted every 1 ps. Each simulation was performed under constant volume conditions at 300 K for 10 ns. For each pH replica, protonation state sampling was enabled throughout the simulation, with Monte Carlo attempts to change protonation states performed every 500 steps. A short relaxation period of 500 steps was applied after each protonation-state change. The replica exchange is attempted every 1 ps between adjacent replicas. All CpHMD and pH-REMD simulations were performed using the explicit solvent implementation^62^ in the AMBER24 software package.^61^ All simulations were performed using the TIP3P water model^41^ with a timestep of 2 fs, with the SHAKE algorithm was enabled.

### QM/MM MD and Umbrella Sampling Simulations

The potential of mean force (PMF) of the proton transfer (PT) between the eDHFR and compound **6** was calculated using QM/MM MD coupled with the umbrella sampling (US) technique.^67^ The QM region comprised the ligand and the side chain of the Asp27 residue (**Figures 5A & S2**), which was treated at the B3LYP-D3/def2-SVP level of theory. The MM region includes the rest of the system, which was treated with the same force field as the classical MD equilibration. The electrostatic embedding scheme was used for the electrostatic interaction between the QM and MM regions. The QM/MM MD simulations were performed at 300 K temperature using a Langevin thermostat and a time step of 0.5 fs. In the US simulation, a collective variable (CV) δ (**Eq 1**) was chosen as the proton transfer coordinate (**Figure 5**) and biased in each umbrella window:

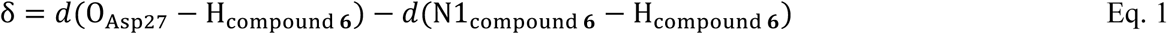

This CV captures the concerted breaking and formation of the N-H and O-H bonds of the donor and acceptor during the PT event. Negative CV value represents the proton being closer to the carboxylic oxygen atom of the Asp27 residue than to the N1 atom on the diaminopyrimidine ring of compound **6**, and *vice versa*. In total, 41 umbrella windows whose harmonic centers were evenly distributed in a range of-1.5 to 1.5 Å of the CV were simulated. For each window, a harmonic potential with a force constant of 50 kcal/mol/Å^2^ was applied and the QM/MM MD trajectory was propagated for approximately 6 ps, removing 1 ps as equilibration, resulting in a total QM/MM US simulation time of approximately 205 ps. The unbiased PMF along the above-defined CV was reconstructed using the weighted histogram analysis method (WHAM)^68^ from the biased CV distributions in each window, and statistical uncertainties were estimated via block average analysis. All QM/MM MD simulations were carried out at B3LYP-D3/def2-SVP/AMBER using TeraChem^56, 55^ interfaced with OpenMM^69^, and the external harmonic bias was applied via interface with the PLUMED plugin.^70^

### Thermodynamic Integration (TI) simulations

TI simulations and thermodynamic cycles were employed to quantify two types of relative binding free energies. The first type is the relative binding free energies between the *cis* and *trans* isomers of each photoswitchable compound, referred to as the *cis*-vs-*trans* affinity differential (ΔΔG_X, 𝑐𝑖𝑠 → 𝑡𝑟𝑎𝑛𝑠_). To calculate this quantity in the TI simulations, *cis* X was converted to *trans* X via an alchemical transformation in which one phenyl ring gradually disappeared from one side of the N=N double bond and reappeared on the opposite side (**Figure 2**), while retaining the same protonation state. The alchemical transformations of the ligand were carried out in both the protein and aqueous solution, yielding the free energy changes ΔG_bound, 𝑐𝑖𝑠 𝑋→ 𝑡𝑟𝑎𝑛𝑠 𝑋_ and ΔG_unbound, 𝑐𝑖𝑠 𝑋→ 𝑡𝑟𝑎𝑛𝑠 𝑋_, respectively. The protonation state of the ligands can differ between the two environments. These values were then combined using a thermodynamic cycle (**Figure 3A, Eq 2**) to obtain the *cis-*vs*-trans* binding affinity difference for compound X (ΔΔG_X, 𝑐𝑖𝑠 → 𝑡𝑟𝑎𝑛𝑠_). TI simulations employing this alchemical pathway and thermodynamic cycle are hereafter referred to as Method A.

**Figure 2.**
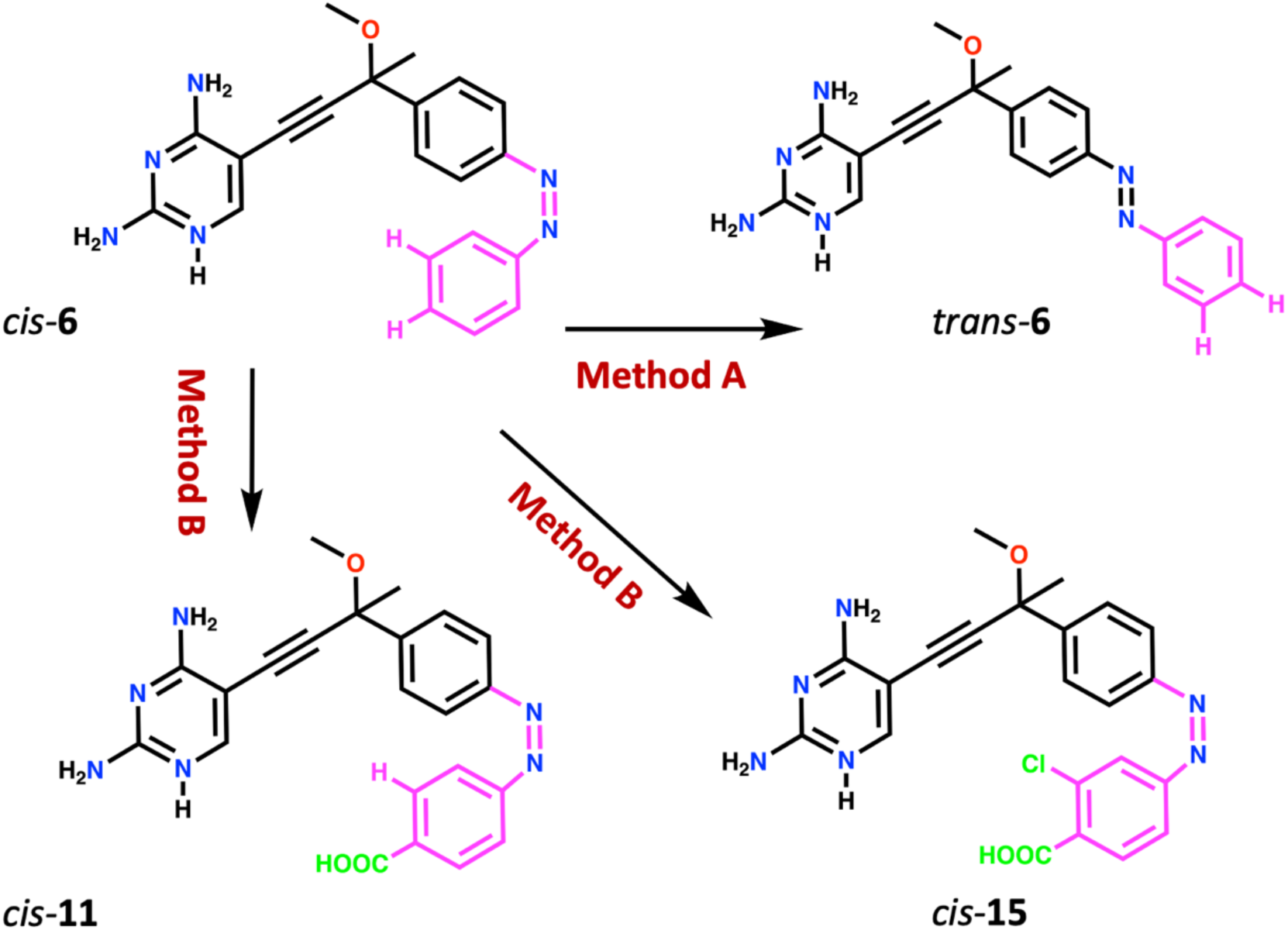
Schematic illustration of the two TI methods for quantifying the *cis*-vs*-trans* affinity differential and substituents’ effects. In Method A, the *cis*-vs*-trans* binding affinity differential is evaluated for a single compound by alchemically transforming from the *cis* to *trans* isomers. This transformation is achieved by relocating a phenyl group across the N=N bond. In Method B, the substituents’ effects on the binding affinity of each isomer are examined by transforming one compound into another while preserving the same isomeric form. In this case, modifications are introduced at specific positions on the benzyl ring, such as para or meta sites. For example, the *cis*-**6** is converted into the *cis*-**11** or *cis*-**15** by replacing a hydrogen atom with a COOH and/or a Cl group at the para and/or meta positions. Both methods can evaluate the substituents’ effects on the *cis*-vs*-trans* binding affinity differential. See Method for details.

**Figure 3.**
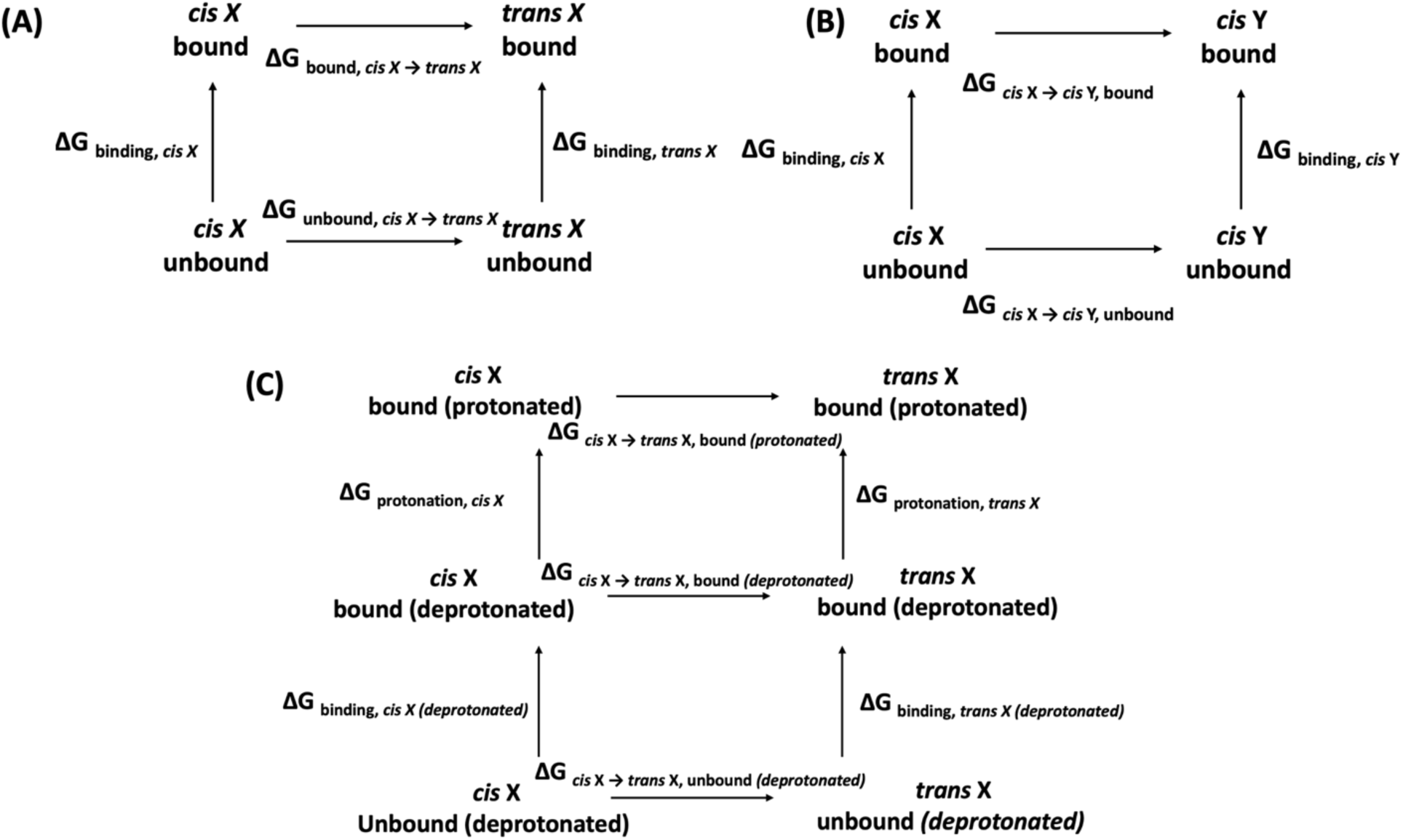
Thermodynamic cycles used for quantifying *cis*-vs-*trans* affinity differential and substituent effects. (A) In Method A, the *cis*-X is converted into *trans*-X through an alchemical transformation performed separately in the protein-bound state (ΔG_bound, 𝑐𝑖𝑠 𝑋→ 𝑡𝑟𝑎𝑛𝑠 𝑋_) and in aqueous solution (ΔG_unbound, 𝑐𝑖𝑠 𝑋→ 𝑡𝑟𝑎𝑛𝑠 𝑋_). The corresponding free energy changes obtained from TI simulations are combined to yield the *cis-*vs*-trans* affinity differential (ΔΔG_X, 𝑐𝑖𝑠 → 𝑡𝑟𝑎𝑛𝑠_) according to Eq 2. (B) In Method B, X is alchemically transformed into Y while maintaining the same isomeric form and protonation state. The associated free energy changes are computed in both the protein (ΔG_𝑐𝑖𝑠 X→𝑐𝑖𝑠 Y, bound_) and aqueous solution (ΔG_𝑐𝑖𝑠 X→𝑐𝑖𝑠 Y, unbound_). These values are combined to calculate the relative binding affinities between X and Y for each isomeric form (ΔΔG_binding, 𝑐𝑖𝑠 X→𝑐𝑖𝑠 Y_, ΔΔG_binding, 𝑡𝑟𝑎𝑛𝑠 X→𝑡𝑟𝑎𝑛𝑠 Y_), according to Eqs. 3 and 4. For both Methods A & B, the combined results from these calculations allow direct evaluation of substituents’ effects on the *cis*-vs-*trans* affinity differential using Eq. 5. (C) For Protocol 1 of Method B, which describes protonation state change of the ligand upon protein binding, the thermodynamic cycle shown in B can be thought of as a combination of two sub-cycles. The upper cycle describes the protonation state change of the ligand in the same environment, while the lower cycle describes the ligand’s binding process in the same protonation state. The overall ΔΔG_binding, 𝑐𝑖𝑠 X→𝑐𝑖𝑠 Y_ and ΔΔG_binding, 𝑡𝑟𝑎𝑛𝑠 X→𝑡𝑟𝑎𝑛𝑠 Y_ calculated using this protocol adds up both the relative pKa’s of the two ligands (upper subsycle, two vertical arrows) and the relative binding affinity between the two ligands in the same protonation states (lower subsycle, two vertical arrows). For Protocol 2 of Method B, only the lower subcycle is involved, and no protonation state change is considered.

The second type of relative binding affinities is between a pair of compounds in the same isomer, yielding ΔΔG_binding, 𝑐𝑖𝑠 X→𝑐𝑖𝑠 Y_ and ΔΔG_binding, 𝑡𝑟𝑎𝑛𝑠 X→𝑡𝑟𝑎𝑛𝑠 Y_. To calculate this quantity, compound X was alchemically transformed into Y by gradually changing the substituent groups that differ between the two molecules, while retaining the isomeric form and protonation state (**Figure 2**). The free energy changes for these transformations were computed in both the protein (ΔG𝑐𝑖𝑠 X→𝑐𝑖𝑠 Y, bound ^and ΔG^𝑡𝑟𝑎𝑛𝑠 X→𝑡𝑟𝑎𝑛𝑠 Y, bound ^) and the aqueous solution^ (ΔG_𝑐𝑖𝑠 X→𝑐𝑖𝑠 Y, unbound_ and ΔG_𝑡𝑟𝑎𝑛𝑠 X→𝑡𝑟𝑎𝑛𝑠 Y, unbound_). The protonation state of the ligands can differ between the two environments. These values were combined using a thermodynamic cycle (**Figure 3B, Eqs 3 & 4**) to predict the relative binding free energies, ΔΔG_binding, 𝑐𝑖𝑠 X→𝑐𝑖𝑠 Y_ and ΔΔG_binding, 𝑡𝑟𝑎𝑛𝑠 X→𝑡𝑟𝑎𝑛𝑠 Y_. TI simulations employing this alchemical pathway and thermodynamic cycle are hereafter referred to as Method B.

Finally, the impact of substituent modification on the *cis*-vs*-trans* affinity difference, denoted as ΔΔΔG_X→Y,𝑐𝑖𝑠–vs–𝑡𝑟𝑎𝑛𝑠 𝑏𝑖𝑛𝑑𝑖𝑛g_, was derived from the relative binding free energies obtained from either method using **Eq 5**. This quantity indicates how the chemical substitutions that convert X to Y modulate the light-responsive affinity differential.

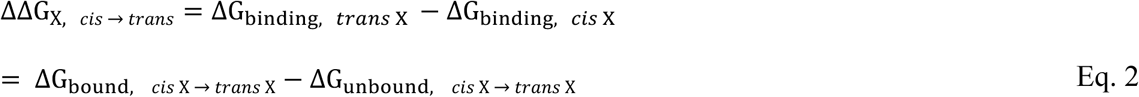

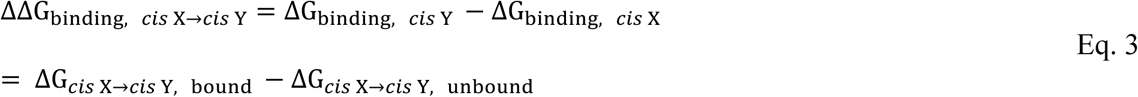

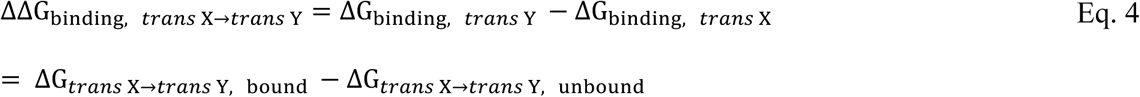

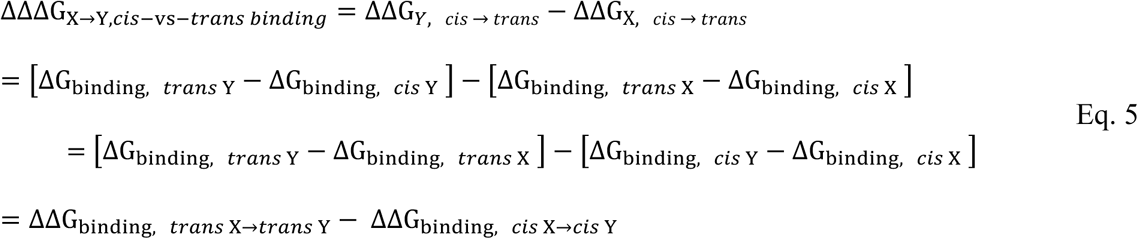

For each alchemical transformation, the atoms disappearing from the initial molecule or appearing in the final one were first defined in a dual topology setup. Then, the free-energy calculation was carried out using a three-stage protocol designed for numerical stability. First, the charges of the disappearing atoms were gradually reduced to zero. Second, the nonbonded and bonded interactions associated with the disappearing atoms were reduced to zero, while those for the appearing atoms were increased to full strength. Both the disappearing and appearing atoms have zero point-charges in this step. Softcore vdW potentials were employed to smooth steric clashes and increase numeric stability. Third, the charges of the appearing atoms were gradually increased to their full values. Each of these three stages was simulated using 11 equally spaced windows (Δλ = 0.1), resulting in a total of 33 λ windows for each transformation.

The starting geometry for each TI simulation was selected from a representative snapshot of the production MD trajectory for the initial compound/isomer X. Then, it is equilibrated using a mixed Hamiltonian of the X and Y (final compound/isomer) connected by the alchemical transformation, at window λ = 0.5. Specifically, with this mixed Hamiltonian, the standard-MD-equilibrated structures were first minimized for 15,000 steps with the steepest descent algorithm, followed by gradual heating to 300 K over 1 ns using a Langevin thermostat^57^ with a collision frequency of 1 ps-1. The systems were then equilibrated for 20 ns in the constant NVT ensemble at 300 K and 1 atm. The final equilibrated structure in the λ = 0.5 mixed Hamiltonian served as the starting point for all λ windows across the three stages. For each λ value in each stage, the system was equilibrated for 500 ps in the constant NVT ensemble, followed by at least 10 ns of production sampling in the constant NPT ensemble. During all TI simulations, bonds involving atoms undergoing alchemical changes were excluded from the SHAKE constraints,^60^ and a 1 fs time step was used consistently throughout. The error bars in the free energy changes were evaluated using block-averaging analysis.

### MM-PBSA and MM-GBSA simulations

Binding free energies were also estimated using the MM-PB(GB)SA methods^34, 35^ based on the final 50 ns of the production MD trajectories. In these calculations, the dielectric constants for the aqueous solution and the protein were set to 80 and 4, respectively. A solvent probe radius of 1.4 Å was used for defining the solvent-accessible surface, and standard force-field atomic radii were applied.^71^ The binding free energies were computed for both isomeric forms of each compound (ΔG_binding, 𝑡𝑟𝑎𝑛𝑠 X_, ΔG_binding, 𝑐𝑖𝑠 X_). Then, the *cis*-vs-*trans* binding affinity differential, ΔΔG_X, 𝑐𝑖𝑠 → 𝑡𝑟𝑎𝑛𝑠_, was obtained from these values using **Eq 2**. The ΔΔΔG_X→Y,𝑐𝑖𝑠–vs–𝑡𝑟𝑎𝑛𝑠 𝑏𝑖𝑛𝑑𝑖𝑛g_ values was subsequently derived according to **Eq 5**.

## Results

This section is organized as follows. First, we investigate the change in the protonation state of the ligands’ diaminopyrimidine ring upon protein binding using pH-REMD and QM/MM umbrella-sampling simulations. Second, we benchmark the performance of TI and endpoint free-energy methods to characterize the *cis*-vs-*trans* affinity differential and the substituents’ effects against the experimental data. The importance of explicitly accounting for the ligand’s binding-induced change in protonation state was highlighted. Also, the accuracy of TI simulations employing two different alchemical transformation pathways was compared and discussed. Third, based on MD simulations, we interpret the protein-ligand interactions that lead to the *cis*-vs-*trans* affinity differential for each ligand under investigation.

### Ligand protonation upon binding to the active site of eDHFR

The diaminopyrimidine ring is shared in all three compounds investigated: compounds **6, 11**, and **15**. To calculate the change in protonation state of the N1 atom on the diaminopyrimidine ring of the three ligands upon binding to the protein, pH-REMD simulations were performed to characterize the pKa’s of compound **6** in both aqueous solution and the protein-bound state. Compound **6** was selected as a representative case of all three compounds, since the N1 atom is well-separated from the additional substituents in compounds **11** and **15** (**Figure 1**), which are unlikely to significantly interfere with the pKa of the diaminopyrimidine ring atom. Based on these simulations, the fraction of the deprotonated state for the ionizable group is plotted as a function of pH, yielding the titration curves (**Figure 4**). These curves were then fitted to the Hill equation to estimate the pKa of the ionizable group.

**Figure 4.**
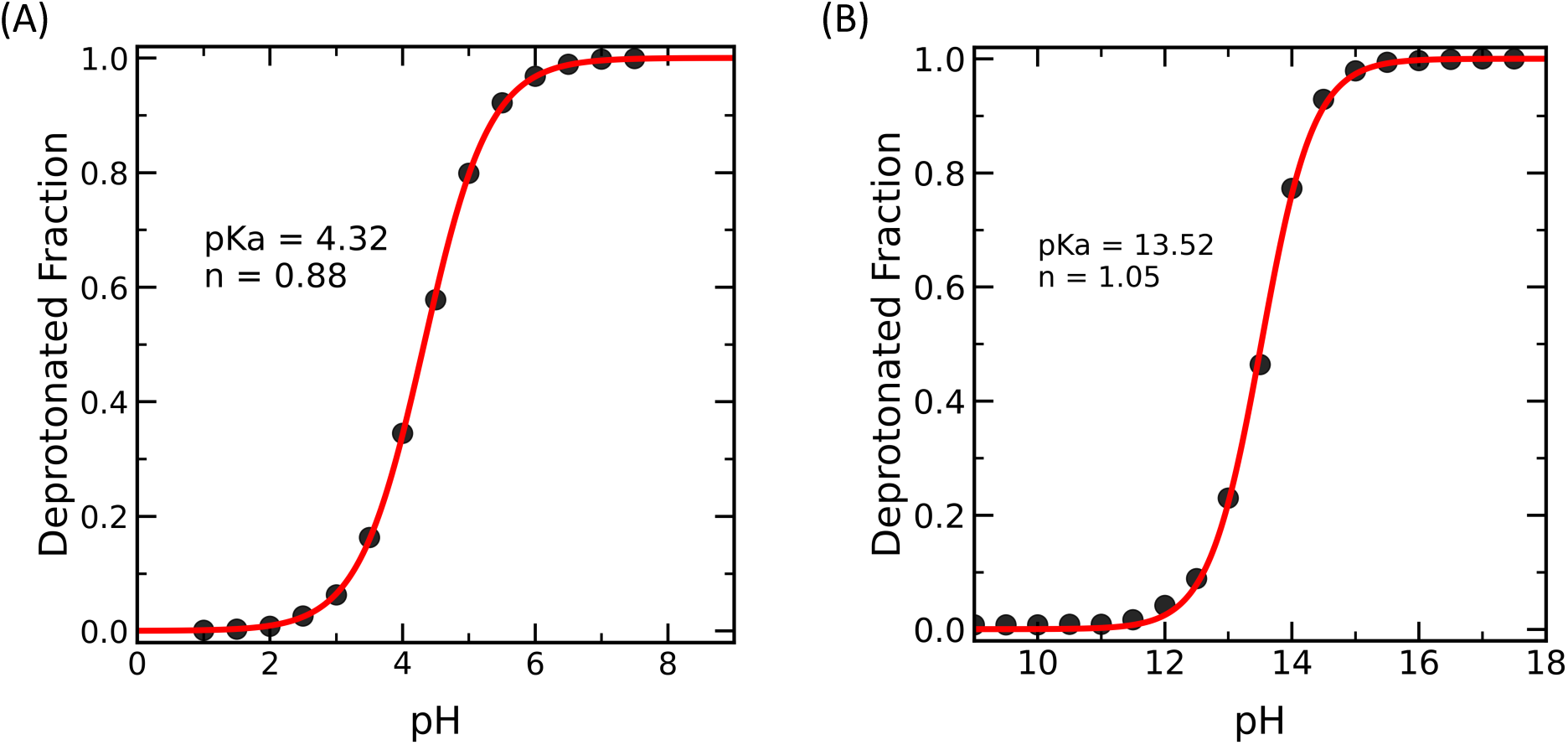
The pH-titration curves of *cis*-**6** obtained from pH-REMD simulations. (A) Aqueous solution system. (B) eDHFR-ligand complex system. The calculated pKₐ values of the ligand substantially increase upon protein binding. In the pH-REMD simulations of the protein-ligand complex, Asp27 was treated as ionizable but remained deprotonated throughout the pH range covered. The Hill coefficients (n’s) are also indicated.

**Figure 5.**
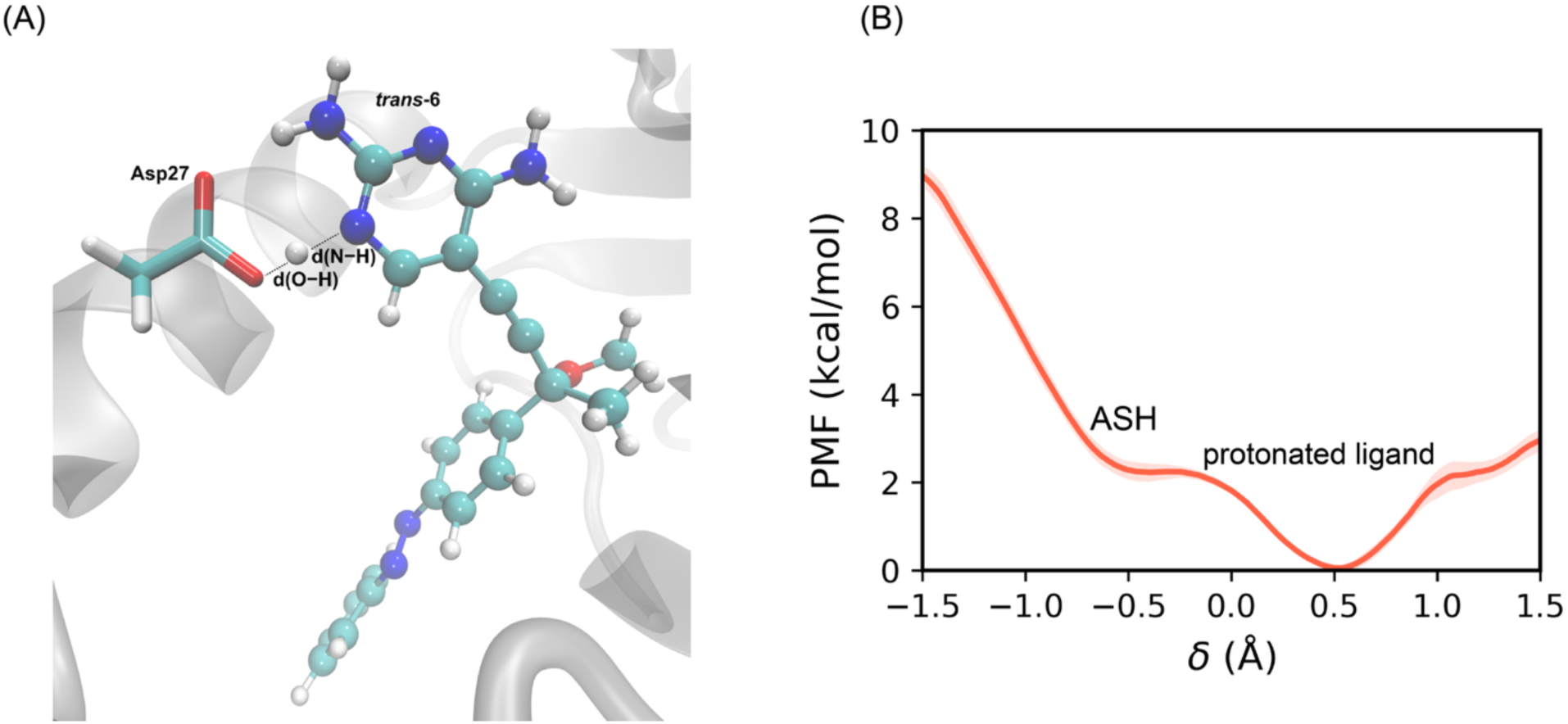
The free energy profile of proton transfer between the Asp27 residue and *trans*-**6**. (A) The salt bridge between the Asp27 and the ligand in the binding site, highlighting the transferring proton and the two key distances for this reaction: d(O–H) and d(N–H). (B) Potential of mean force (PMF) obtained from B3LYP-D/MM US simulations along the reaction coordinate δ = d(O– H) − d(N–H). The free energy profile shows a lowest minimum in the region where the proton resides on the ligand (δ = ∼ 0.5Å), indicating that the protonated ligand/deprotonated Asp27 is thermodynamically more favored than the deprotonated ligand/protonated Asp27. The shaded belt represents the error bars.

In the pH-REMD simulations, the reference pKa value in aqueous solution was set to 4.3 for compound **6**. This reference pKa value was calculated using M06-2X/TZVP/CPCM (𝜀 = 80.15, see Method for details).^64^ It is worth noting that the TMP, which served as the parent compound for the design of compounds **6**, has an experimental pKa of 7.12 in bulk solution.^72^ Using the same electronic structure method, the calculated pKa of TMP is 8.61. Thus, our DFT calculation predicts that the pKa of compound **6** is ∼4.3 units lower than TMP. Although this pKa shift could be overestimated, it suggests that the chemical modifications introduced in TMP reduce its basicity, likely due to changes in local charge distribution. Considering that the experimental pKa of 7.12 for the TMP^72^ and our DFT calculated pKa shift, the compound **6** is most likely deprotonated in the aqueous solution in near-neutral pH conditions.

The pKa of the N1 atom in compound **6,** calculated by pH-REMD simulation, is increased to 13.5 from 4.3 upon binding to the active site of eDHFR (**Figure 4B**). Importantly, in the pH-REMD simulations of the protein-ligand complex, Asp27 was treated as ionizable but remained predominantly deprotonated (>97% of the population) throughout the covered pH range. This ∼9 pH unit upward shift in the ligand’s pKa indicates strong stabilization of the ligand’s protonated state within the protein environment, mostly due to forming a hydrogen bond with the deprotonated Asp27 residue. As a result, the pH-titration curve confirms that compound **6** remains predominantly N1-protonated near physiological pH range.

The large shift in the ligand’s pKa upon protein binding highlights the strong influence of the local electrostatic environment on the ligand’s protonation equilibrium. In this case, strong hydrogen-bond interactions with Asp27 favor the protonation of the N1 atom. Importantly, these pH-REMD simulation results are consistent with the proton transfer free energy profile obtained from the QM/MM umbrella sampling simulations (see below), which also indicate that the protonated form of the diaminopyrimidine ring is stabilized in the protein-bound state. Furthermore, the results align with previous experimental studies, in which the diaminopyrimidine rings of the TMP and methotrexate in complex with eDHFR have pKa values above 10 and are considered protonated.

To further validate the results from constant pH simulations, we performed QM/MM US simulations to characterize the potential of mean force (PMF) of proton transfer between the N1 atom of the compound **6** and the carboxylic group on the Asp27 residue in the active site of eDHFR (Figures 5A). The QM region includes the side chain of Asp27 and the whole ligand, which was treated at the B3LYP-D3/def2-SVP level of theory (Method). The PMF is constructed along the proton transfer coordinate (δ) (Figure 5A and **Eq 1**). The PMF (Figure 5B) features a lowest-energy minimum at δ = ∼ 0.5 Å, corresponding to the Asp27-deprotonated/N1-protonated state. Meanwhile, a shoulder in the PMF at δ = ∼-0.5 Å corresponding to the Asp27-protonated/N1-deprotonated state has more than 2 kcal/mol higher free energy than the minimum at δ = ∼ 0.5 Å. Thus, the PMF demonstrates that proton transfer from Asp27 to the ligand is thermodynamically favorable in the protein environment. In other words, when the Asp27’s carboxylic group and the ligand’s N1 atom form a hydrogen bond, the proton involved in this hydrogen bond prefers to be localized on the ligand. The higher proton binding affinity of the ligand than that of Asp27 likely arises from the delocalization of the positive charge over the diaminopyrimidine ring, which helps stabilize its protonated state. The positively charged, N1-protonated diaminopyrimidine ring can thus form a strong salt bridge with the negatively charged, deprotonated Asp27. Removal of the proton from the ligand not only needs to overcome the intrinsic proton affinity of the diaminopyrimidine ring in isolation, but also break the strong salt-bridge interaction between the ligand and the Asp27. Therefore, this salt-bridge interaction further stabilizes the ligand’s protonated state, greatly increasing its pKa in the active site of eDHFR. This observation from the *ab initio* QM/MM PMF calculation is consistent with our MM-level pH-REMD simulations. Importantly, the characterization of the ligand’s protonation state has direct implications for the interpretation of binding free energy calculations discussed below. Without explicit consideration of the N1-atom’s protonation state change upon protein binding, no free energy method tested could consistently predict the experimentally observed *cis*-vs-*trans* affinity differential or the substituents’ effects on it.

### Method Benchmark: cis-vs-trans affinity differential and substituents’ effects

The experimental design campaign of the TMP-derived photoswitchable eDHFR inhibitor^17^ has shown that chemical modification can substantially alter the absolute and light-responsive affinity differential of these ligands. These substitutions were introduced at the *para* and *meta* positions of the phenyl ring in the azobenzene moiety. In particular, introducing a carboxylic acid group to compound **6**, yielding compound **11**, increases the ligand’s inhibitory potency but eliminates the “*cis*-on” effect. Meanwhile, introducing an additional chlorine atom on the azobenzene moiety of compound **11**, yielding compound **15**, recovers the “*cis*-on” effect.

These prior experimental results provide a valuable dataset for benchmarking free energy methods. Here, we evaluated the accuracy of TI and MM-PB/GBSA methods in predicting the *cis*-vs-*trans* affinity differential and its substituents’ effects. The experimental reference dataset consists of relative binding affinities between different ligands (compounds **6, 11**, and **15**) in the same isomeric form, as well as the *cis*-vs-*trans* affinity differential for each ligand. They were derived from the ratios of reported IC_50_ values.^17^

### 1. TI simulations with different alchemical transformation pathways

Given its established accuracy for photoswitchable ligands targeting G-protein coupled receptor and tubulin^18, 19^, the TI method was first tested. Due to finite sampling, the difficulty of converging the free energy changes often depends on the selected thermodynamic cycle and alchemical transformation pathways. Here, we compared two pathways. In the first pathway (Method A), the *cis* and *trans* isomers are interconverted by gradually making a phenyl ring attached to the N=N double bond disappear from one side and reappear on the other. Such an alchemical transformation was performed in both the aqueous solution and the eDHFR binding pocket. Based on the above-described pKa calculations, the N1 atom of all three ligands was first chosen to be deprotonated in the aqueous solution and protonated in the protein. This approach yields the *cis*-vs-*trans* affinity differential for each compound X through a thermodynamic cycle (Figure 3A, **Eq 2**), i.e.,ΔΔG_X, 𝑐𝑖𝑠 → 𝑡𝑟𝑎𝑛𝑠_. By taking the difference of this quantity for a pair of compounds, X and Y, the substituent’s effect on the *cis*-vs-*trans* affinity differential, ΔΔΔG_X→Y, 𝑐𝑖𝑠–vs–𝑡𝑟𝑎𝑛𝑠 𝑏𝑖𝑛𝑑𝑖𝑛g_, can be obtained. *Accurately predicting this property is key to rational design aimed at increasing the ligand’s light responsiveness via chemical substitution.* A positive value of this quantity indicates that changing the substituents from X to Y enhances the “*cis*-on” effect. The resulting ΔΔG_X, 𝑐𝑖𝑠 → 𝑡𝑟𝑎𝑛𝑠_ and ΔΔΔG_X→Y, 𝑐𝑖𝑠–vs–𝑡𝑟𝑎𝑛𝑠 𝑏𝑖𝑛𝑑𝑖𝑛g_ obtained from Method A are summarized alongside experimental data in **Tables 1 & 3**.

**Table 1.** Calculated *cis*-vs-*trans* affinity differentials (ΔΔG_X, 𝑐𝑖𝑠 → 𝑡𝑟𝑎𝑛𝑠_, in kcal/mol) for compounds **6, 11**, and **15**, calculated using TI (Method A) and MM-PB(GB)SA. Experimental values^17^ are included for comparison. Positive ΔΔG_X, 𝑐𝑖𝑠 → 𝑡𝑟𝑎𝑛𝑠_ values indicate “*cis*-on*”* effect, and vice versa. In these calculations, the ligands are N1-deprotonated and N1-protonated in the aqueous solution and protein, respectively.

| Quantity | Compounds |  |  |
| --- | --- | --- | --- |
|  | 6 | 11 | 15 |
| $\Delta\Delta G_{X, \text{ cis} \rightarrow \text{trans}}$ (Exp.) | $0.5 \pm 0.5$ | $-0.1 \pm 0.4$ | $0.4 \pm 0.2$ |
| $\Delta G_{\text{unbound}, \text{ cis } X \rightarrow \text{trans } X}$ (TI) | $-11.9 \pm 0.1$ | $-11.9 \pm 0.1$ | $-11.6 \pm 0.1$ |
| $\Delta G_{\text{bound}, \text{ cis } X \rightarrow \text{trans } X}$ (TI) | $-11.3 \pm 0.5$ | $-13.0 \pm 0.1$ | $-11.3 \pm 0.1$ |
| $\Delta\Delta G_{X, \text{ cis} \rightarrow \text{trans}}$ (TI) | $0.6 \pm 0.5$ | $-1.1 \pm 0.1$ | $0.3 \pm 0.1$ |
| $\Delta G_{\text{binding}, \text{ cis } X}$ (MM-GBSA) | $-64.7 \pm 0.1$ | $-72.0 \pm 0.1$ | $-73.6 \pm 0.1$ |
| $\Delta G_{\text{binding}, \text{ trans } X}$ (MM-GBSA) | $-57.9 \pm 0.1$ | $-56.5 \pm 0.1$ | $-54.9 \pm 0.1$ |
| $\Delta\Delta G_{X, \text{ cis} \rightarrow \text{trans}}$ (MM-GBSA) | $6.8 \pm 0.1$ | $15.5 \pm 0.1$ | $18.7 \pm 0.1$ |
| $\Delta G_{\text{binding}, \text{ cis } X}$ (MM-PBSA) | $-35.5 \pm 0.1$ | $-33.7 \pm 0.1$ | $-34.6 \pm 0.1$ |
| $\Delta G_{\text{binding}, \text{ trans } X}$ (MM-PBSA) | $-27.0 \pm 0.1$ | $-29.9 \pm 0.1$ | $-32.1 \pm 0.1$ |
| $\Delta\Delta G_{X, \text{ cis} \rightarrow \text{trans}}$ (MM-PBSA) | $8.5 \pm 0.1$ | $3.8 \pm 0.1$ | $2.5 \pm 0.1$ |

**Table 2.** Calculated binding free energy changes (ΔΔG _binding, X→Y_, in kcal/mol) upon converting compound X to Y, calculated using TI (Method B). Experimental values^17^ are included for comparison. Negative values of ΔΔG _binding, X→Y_ correspond to an increase in binding affinity upon converting X to Y, and *vice versa*. Calculations were performed using two different protocols. In Protocol 1, the ligands were in the N1-protonated state in the protein and the N1-deprotonated state in the aqueous solution. In Protocol 2, the ligands were in the same protonation state (either N1-protonated or N1-deprotonated) in both environments.

| Transformations | Quantities |  |  |  |
| --- | --- | --- | --- | --- |
| | $\Delta G_{\text{X}\rightarrow\text{Y, unbound}}$ | $\Delta G_{\text{X}\rightarrow\text{Y, bound}}$ | $\Delta\Delta G_{\text{binding, X}\rightarrow\text{Y}}$<br>(TI) | $\Delta\Delta G_{\text{binding, X}\rightarrow\text{Y}}$<br>(Exp.) |
| **Protocol 1** |  |  |  |  |
| **11→15**<br>(*cis*) | -26.8 ± 0.5 | -4.2 ± 0.6 | 22.6 ± 0.8 | -0.6 ± 0.3 |
| **11→15**<br>(*trans*) | -27.2 ± 0.3 | -4.3 ± 0.5 | 22.9 ± 0.6 | -0.1 ± 0.2 |
| **6→11**<br>(*cis*) | -57.8 ± 0.0 | -8.2 ± 0.0 | 49.6 ± 0.0 | -0.5 ± 0.4 |
| **6→11**<br>(*trans*) | -57.9 ± 0.1 | -5.8 ± 0.0 | 52.1 ± 0.1 | -1.1 ± 0.3 |
| **Protocol 2** |  |  |  |  |
| **N1-protonated** |  |  |  |  |
| **11→15**<br>(*cis*) | -3.1 ± 0.2 | -4.2 ± 0.6 | -1.1 ± 0.6 | -0.6 ± 0.3 |
| **11→15**<br>(*trans*) | -3.8 ± 0.4 | -4.3 ± 0.5 | -0.5 ± 0.6 | -0.1 ± 0.2 |
| **6→11**<br>(*cis*) | -7.6 ± 0.0 | -8.2 ± 0.0 | -0.6 ± 0.0 | -0.5 ± 0.4 |
| **6→11**<br>(*trans*) | -7.4 ± 0.1 | -5.8 ± 0.0 | 1.6 ± 0.1 | -1.1 ± 0.3 |
| **N1-deprotonated** |  |  |  |  |
| **11→15**<br>(*cis*) | -26.8 ± 0.5 | -27.0 ± 0.7 | -0.2 ± 0.9 | -0.6 ± 0.3 |
| **11→15**<br>(*trans*) | -27.2 ± 0.3 | -26.4 ± 0.4 | 0.8 ± 0.5 | -0.1 ± 0.2 |
| **6→11**<br>(*cis*) | -57.8 ± 0.0 | -55.4 ± 0.5 | 2.4 ± 0.5 | -0.5 ± 0.4 |
| **6→11**<br>(*trans*) | -57.9 ± 0.1 | -55.8 ± 0.1 | 2.1 ± 0.1 | -1.1 ± 0.3 |

**Table 3.**
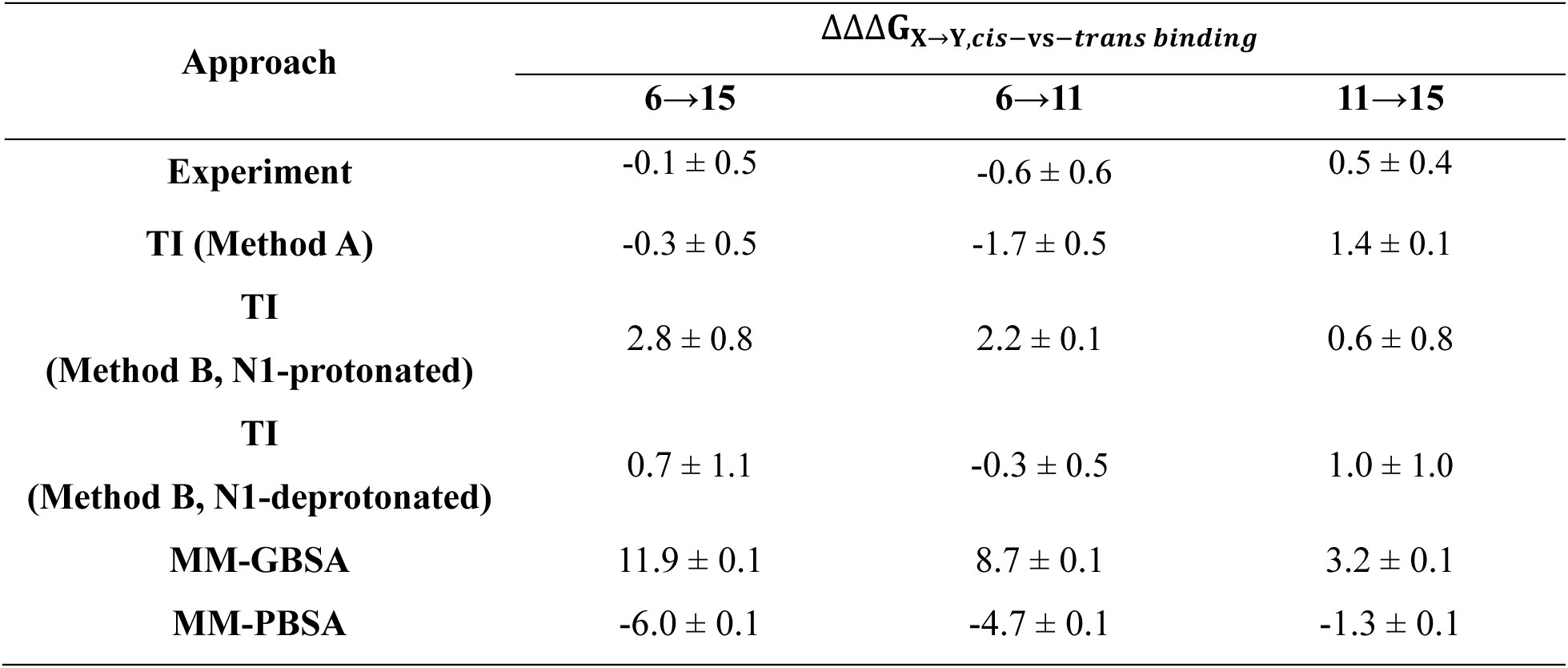
Substituents’ effects in light-responsive affinity differential (ΔΔΔG _X→Y,𝑐𝑖𝑠–vs–𝑡𝑟𝑎𝑛𝑠 𝑏𝑖𝑛𝑑𝑖𝑛g_ in kcal/mol) upon converting X to Y, evaluated using TI (Methods A and B) and MM-PB(GB)SA simulations, and compared with values.^17^ Positive values indicate that X→Y conversion enhances the “*cis*-on” effects, and *vice versa*.

In the second pathway (Method B), compound X is converted to Y in the same isomeric form (*cis* or *trans*) in both the aqueous solution and the protein-ligand complex. Through a thermodynamic cycle (Figure 2B), this approach estimates the relative binding free energy between the two compounds in identical isomeric form and protonation state, i.e., ΔΔG_binding, 𝑐𝑖𝑠 X→𝑐𝑖𝑠 Y_ and ΔΔG_binding, 𝑡𝑟𝑎𝑛𝑠 X→𝑡𝑟𝑎𝑛𝑠 Y_ (Figure 2B and **Eqs. 3 &4**). The difference between these two ΔΔG_binding_ values corresponds to the substituent’s effect on the *cis*-vs-*trans* affinity differential, ΔΔΔG_X→Y, 𝑐𝑖𝑠–vs–𝑡𝑟𝑎𝑛𝑠 𝑏𝑖𝑛𝑑𝑖𝑛g_, which is also evaluated by Method A. The ligand’s protonation state in solution and in the protein can be the same or different. **Tables 2 & 3** summarize the results obtained from this approach with the same protonation state in solution and protein (reasons for this choice are discussed below), and experimental values. In our previous studies, this method has provided the best quantitative interpretation of experimentally observed trends for a series of photoswitchable ligands targeting tubulin and membrane receptors.^18, 19^

Using Method A, the *cis*-vs-*trans* affinity differentials (ΔΔG_X, 𝑐𝑖𝑠 → 𝑡𝑟𝑎𝑛𝑠_) for all three compounds **6, 11** and **15** reproduced the experiment with near-quantitative accuracy (**Table 1**). The TI simulations correctly predicted the signs of *cis*-vs-*trans* affinity differentials (ΔΔG_X, 𝑐𝑖𝑠 → 𝑡𝑟𝑎𝑛𝑠_), i.e., the “*cis*-on” or “*trans*-on” effects. The positive ΔΔG_X, 𝑐𝑖𝑠 → 𝑡𝑟𝑎𝑛𝑠_ values of compounds **6** and **15** indicate “*cis*-on” effect, whereas the small negative ΔΔG_X, 𝑐𝑖𝑠 → 𝑡𝑟𝑎𝑛𝑠_ of compound **11** displays indicates a slight “*trans*-on” effect. Beyond this, Method A also performs well in predicting how chemical substitution alters the *cis*-vs*-trans* affinity differential (**Table 3**). The calculated ΔΔΔG_X→Y,𝑐𝑖𝑠–vs–𝑡𝑟𝑎𝑛𝑠 𝑏𝑖𝑛𝑑𝑖𝑛g_ values indicate that the **6**→**11** transformation (introducing a carboxylate group) neutralizes the “*cis*-on” effect and results in a slight “*trans*-on” effect. In contrast, the **11**→**15** transformation (introducing a chlorine atom on top of the carboxylate group) recovers the “*cis*-on” effect of compound **6**. These calculated results are all consistent with experiments, indicating that Method A is a robust framework for predicting both active isomer and substituents’ effects in their light-responsive affinity differential. The ability of Method A to quantitatively predict whether a ligand is *cis*-on or *trans*-on, and how chemical modifications magnify, mitigate, or reverse these effects, is critical for rational design of photoswitchable ligands in photopharmacology.

Method A does not perform as well if the protonation states are set the same in both the solution and aqueous environment, as shown in **Table S1**. Specifically, when the ligand is set as N1-deprotonated in both environments, the calculated ΔΔG_X, 𝑐𝑖𝑠 → 𝑡𝑟𝑎𝑛𝑠_ for compounds 11 and 15 are largely overestimated, and the ΔΔΔG_X→Y,𝑐𝑖𝑠–vs–𝑡𝑟𝑎𝑛𝑠 𝑏𝑖𝑛𝑑𝑖𝑛g_ values fail to predict the sign of experimental data for any of the three transformations, generating the worst prediction. When the ligand is set as N1-protonated in both environments, the calculated ΔΔG_X, 𝑐𝑖𝑠 → 𝑡𝑟𝑎𝑛𝑠_ for compound 11 is largely underestimated, so is the ΔΔΔG_X→Y,𝑐𝑖𝑠–vs–𝑡𝑟𝑎𝑛𝑠 𝑏𝑖𝑛𝑑𝑖𝑛g_ for 6→11 transformation. Also, the calculated ΔΔΔG_X→Y,𝑐𝑖𝑠–vs–𝑡𝑟𝑎𝑛𝑠 𝑏𝑖𝑛𝑑𝑖𝑛g_ for the 6→15 has the incorrect sign. These negative results highlight that the binding-induced changes in the ligand’s protonation state must be explicitly accounted for in TI simulations to obtain accurate predictions of light-responsive affinity differential and substituents’ effects.

Next, we benchmarked the accuracy of Method B. This method uses the free energy change of alchemical transformation from X to Y in the protein and aqueous solution (e.g., ΔG_𝑡𝑟𝑎𝑛𝑠 X→𝑡𝑟𝑎𝑛𝑠 Y, bound_ and ΔG_𝑡𝑟𝑎𝑛𝑠 X→𝑡𝑟𝑎𝑛𝑠 Y, unbound_) to estimate the relative binding affinity between a pair of compounds (e.g., ΔΔG_binding, 𝑡𝑟𝑎𝑛𝑠 X→𝑡𝑟𝑎𝑛𝑠 Y_). In principle, one should simulate the alchemical transformation in the protein-bound state (e.g., ΔG_𝑡𝑟𝑎𝑛𝑠 X→𝑡𝑟𝑎𝑛𝑠 Y, bound_) with N1-protonated X and Y, and that in the aqueous solution state (e.g., ΔG_𝑡𝑟𝑎𝑛𝑠 X→𝑡𝑟𝑎𝑛𝑠 Y, unbound_) with N1-deprotonated X and Y (Protocol 1). Such a choice would, ideally, reflect the binding-induced change in protonation state. Alternatively, one could make a suboptimal choice of setting the ligands to the same protonation state (either N1-protonated or N1-deprotonated) in both the solution and the protein (Protocol 2), which could introduce systematic errors in and ΔΔG_binding, 𝑐𝑖𝑠 X→𝑐𝑖𝑠 Y_. Note that binding-induced changes in the ligand’s protonation state were naturally incorporated in the experimental data. Comparing the performance of Protocols 1 and 2 enabling a systematic analysis of how different treatments of the ligand’s protonation state affect the accuracy of free energy calculations.

The calculated ΔΔG_binding, 𝑡𝑟𝑎𝑛𝑠 X→𝑡𝑟𝑎𝑛𝑠 Y_ and ΔΔG_binding, 𝑐𝑖𝑠 X→𝑐𝑖𝑠 Y_ calculated using Protocol 2 are reported in the **Table 2**, and the *cis*-vs*-trans* affinity differential ΔΔΔG_X→Y,𝑐𝑖𝑠–vs–𝑡𝑟𝑎𝑛𝑠 𝑏𝑖𝑛𝑑𝑖𝑛g_ in **Table 3**. For the **11**→**15** transformation in the N1-protonated state, Method B captures the experimental trend reasonably well. The calculated ΔΔG_binding, 𝑐𝑖𝑠 X→𝑐𝑖𝑠 Y_ (-1.1 ± 0.6 kcal/mol) indicates that introducing an additional chlorine substituent alongside the carboxylate group strengthens the binding affinity of *cis* **15** relative to *cis* **11**, in agreement with the experimental value of-0.6 ± 0.3 kcal/mol. A similar stabilization is observed for the *trans* isomer, where both simulation (-0.5 ± 0.6 kcal/mol) and experiment (-0.1 ± 0.2 kcal/mol) suggest a slight increase in affinity upon conversion from *trans* **11** to *trans* **15**. These results indicate that, for this specific modification, Method B reproduces the qualitative effect of substitution on binding in both isomeric states. For the *cis* **6**→ *cis* **11** transformation in the protonated system in the N1-protonated state, Method B predicts stronger binding upon introducing the carboxylic acid group for the *cis* isomer, in line with experimental data. This is mainly due to the formation of a favorable salt-bridge interaction between the negatively charged carboxylate on *cis* **11** and the Arg57 residue in eDHFR, which is absent in *cis* **6**. However, the performance of Method B deteriorates for the *trans* **6** → *trans* **11** transformation. The experimental data indicate an increase in the binding affinity after this transformation (ΔΔG_binding, 𝑡𝑟𝑎𝑛𝑠 X→𝑡𝑟𝑎𝑛𝑠 Y_ =-1.1 ± 0.3 kcal/mol), whereas the calculation indicates a decrease in the binding affinity (1.6 ± 0.1 kcal/mol).

For the N1-deprotonated state of the ligands, Method B cannot consistently capture the experimental trend. For the *trans* **11**→ *trans* **15** transformation, the calculations predict a decrease in the binding affinity (ΔΔG_binding, 𝑡𝑟𝑎𝑛𝑠 X→𝑡𝑟𝑎𝑛𝑠 Y_ = 0.8 ± 0.5 kcal/mol). This result is qualitatively inconsistent with the experimental value (-0.1 ± 0.2 kcal/mol). A similar loss of accuracy is observed for the **6**→**11** transformation, where the calculations predict a decrease in binding affinity for both isomers, in contrast with the increase in binding affinity observed from experimental data (**Table 2**). The limitations of Method B become more obvious for ΔΔΔG_X→Y,𝑐𝑖𝑠–vs–𝑡𝑟𝑎𝑛𝑠 𝑏𝑖𝑛𝑑𝑖𝑛g_ (**Table 3**). For the **11**→**15** transformation, Method B correctly predicts an enhancement of the “*cis*-on” effect for both protonated and deprotonated ligands, consistent with experiment. However, for the **6**→**11** transformation, Method B fails to reproduce the negative ΔΔΔG_X→Y,𝑐𝑖𝑠–vs–𝑡𝑟𝑎𝑛𝑠 𝑏𝑖𝑛𝑑𝑖𝑛g_ from the experiments. This sign error indicates that Method B cannot reliably resolve the delicate shift in the *cis*-vs-*trans* affinity differential for this transformation. Since Method B uses the difference between ΔΔG_binding, 𝑐𝑖𝑠 X→𝑐𝑖𝑠 Y_ and ΔΔG_binding, 𝑡𝑟𝑎𝑛𝑠 X→𝑡𝑟𝑎𝑛𝑠 Y_ to calculate ΔΔΔG_X→Y,𝑐𝑖𝑠–vs–𝑡𝑟𝑎𝑛𝑠 𝑏𝑖𝑛𝑑𝑖𝑛g_, small inaccuracies in either of the two ΔΔG_binding_ quantities, as discussed above (**Table 2**), can propagate and lead to a qualitatively incorrect prediction of ΔΔΔG_X→Y,𝑐𝑖𝑠–vs–𝑡𝑟𝑎𝑛𝑠 𝑏𝑖𝑛𝑑𝑖𝑛g_.

It is worth noting that a source of systematic error in Protocol 2’s estimates of ΔΔG_binding, 𝑡𝑟𝑎𝑛𝑠 X→𝑡𝑟𝑎𝑛𝑠 Y_ and ΔΔG_binding, 𝑐𝑖𝑠 X→𝑐𝑖𝑠 Y_ is the erroneous assumption that the ligand’s protonation state remains the same in both aqueous solution and protein. However, in practice, the suboptimal Protocol 2 had to be chosen deliberately because of the poor results from the conceptually optimal Protocol 1. When choosing Protocol 1, the ΔΔG_binding, 𝑡𝑟𝑎𝑛𝑠 X→𝑡𝑟𝑎𝑛𝑠 Y_ and ΔΔG_binding, 𝑐𝑖𝑠 X→𝑐𝑖𝑠 Y_ deviate much more from the experiment than Protocol 2 (**Table 2**), often by 1-2 orders of magnitude. For example, the ΔΔG_binding, 𝑐𝑖𝑠 X→𝑐𝑖𝑠 Y_ for *cis* **6**→ *cis* **11** transformation estimated by Protocol 1 is ∼50 kcal/mol, in sharp contrast to the experimental value of-0.5 ± 0.4 kcal/mol and the Protocol 2’s estimated values of-0.6 ± 0.0 kcal/mol (protonated state) and 2.4 ± 0.5 kcal/mol (deprotonated state). The reason for the poor performance of Protocol 1 is exactly what makes it conceptually correct: ligands change protonation state upon binding with the protein. When calculating the relative binding free energies using Protocol 1, its associated thermodynamic cycle (Figure 3C) entails solving two difficult tasks together: (1) estimating relative binding free energies in the same protonation state between X and Y, (2) estimating the relative pKa between X and Y. For task 2, when the force field parameters are not accurately parameterized for this specific purpose, small perturbations in the fitted point charges in both protonation states can readily lead to large errors in the predicted pKa without *ad hoc* reference correction (See Discussion). Although these pKa values are not explicitly calculated for a pair of compounds connected by alchemical transformation, their large errors must cancel out each other to predict correct relative binding free energies. However, such error cancellations are *not* guaranteed, leading to large errors in the ΔΔG_binding, 𝑡𝑟𝑎𝑛𝑠 X→𝑡𝑟𝑎𝑛𝑠 Y_ and ΔΔG_binding, 𝑐𝑖𝑠 X→𝑐𝑖𝑠 Y_ estimates. As a result, the prediction of ΔΔΔG_X→Y,𝑐𝑖𝑠–vs–𝑡𝑟𝑎𝑛𝑠 𝑏𝑖𝑛𝑑𝑖𝑛g_, which is defined as the difference between these two quantities, becomes very unreliable.

In contrast, Protocol 2 involves only task 1 (Figure 3B). For this reason, the best performance of Method B was achieved with Protocol 2, i.e., setting the same protonation state for each ligand in both the protein and the aqueous solution. This approach removes the dominant source of error arising from pKa differences and focuses on the effects of chemical substitution on binding affinity in the same protonation state. While this treatment does not explicitly capture the coupling between protonation and binding, it provides a more physically interpretable estimate of substituent effects within the limitations of non-reactive force fields. It is worth noting that in our previous studies^18, 19^, Method B was identified as the most accurate method for estimating substituents’ effects on light-responsive affinity differentials of photoswitchable inhibitors of tubulin and β-adrenergic receptors. Here, the new comparative analysis highlights a critical limitation of Method B when applied to ionizable, photoswitchable ligands involving changes in protonation state upon protein binding. Method A, when choosing the correct protonation states in each environment, outperforms Method B.

### 2. MM-PBSA and MM-GBSA

The *cis*-vs-*trans* affinity differentials (ΔΔG_X, 𝑐𝑖𝑠 → 𝑡𝑟𝑎𝑛𝑠_) obtained from MM-PB(GB)SA calculations are summarized in **Table 1**. The magnitude of the calculated values deviates significantly from the experimental values for all compounds. Additionally, both methods consistently predict a “*cis-*on” effect for all three compounds. However, this is inconsistent with experimental observations for **11**, which has *a “trans*-on” effect. This indicates that MM-GBSA is not able to consistently identify “*cis*-on” or “*trans*-on” effects correctly.

The limitations of these methods become even more obvious for ΔΔΔG _X→Y,𝑐𝑖𝑠–vs–𝑡𝑟𝑎𝑛𝑠 𝑏𝑖𝑛𝑑𝑖𝑛g_ (**Table 3**). MM-GBSA predicts large positive ΔΔΔG _X→Y,𝑐𝑖𝑠–vs–𝑡𝑟𝑎𝑛𝑠 𝑏𝑖𝑛𝑑𝑖𝑛g_ values for all three transformations (**6**→**11, 6**→**15**, and **11**→**15**), indicating a strong enhancement of “*cis*-on” effect upon these transformations. The magnitudes of these values are an order of magnitude higher than the experimental values, and the signs of them are both wrong (**Table 3**). These results indicate that, even after error cancellation by subtracting the ΔΔG_X, 𝑐𝑖𝑠 → 𝑡𝑟𝑎𝑛𝑠_ values, MM-GBSA still cannot qualitatively predict substituent effects on the differential light-responsive affinity. MM-PBSA, on the other hand, predicts negative ΔΔΔG _X→Y,𝑐𝑖𝑠–vs–𝑡𝑟𝑎𝑛𝑠 𝑏𝑖𝑛𝑑𝑖𝑛g_ values for all three transformations, suggesting that they reduce the “*cis*-on” effect. While this qualitative trend partially agrees with the experiment for **6**→**11** and **6**→**15**, it is inconsistent for **11**→**15**, where the experiment indicates an increase in “*cis-*on” effect. Overall, these end-point free energy methods cannot qualitatively predict the *cis-*vs*-trans* affinity differentials or substituent effects.

### Protein-ligand interactions in eDHFR

To understand the molecular basis of the *cis*-vs-*trans* affinity differentials of the TMP derivatives, we analyzed electrostatic and van der Waals (vdW) interaction energies between the ligands and protein, as well as between the ligands and surrounding water molecules, using production MD trajectories (Figure 6). We also compared representative binding poses in the eDHFR active site (Figure 7).

**Figure 6.**
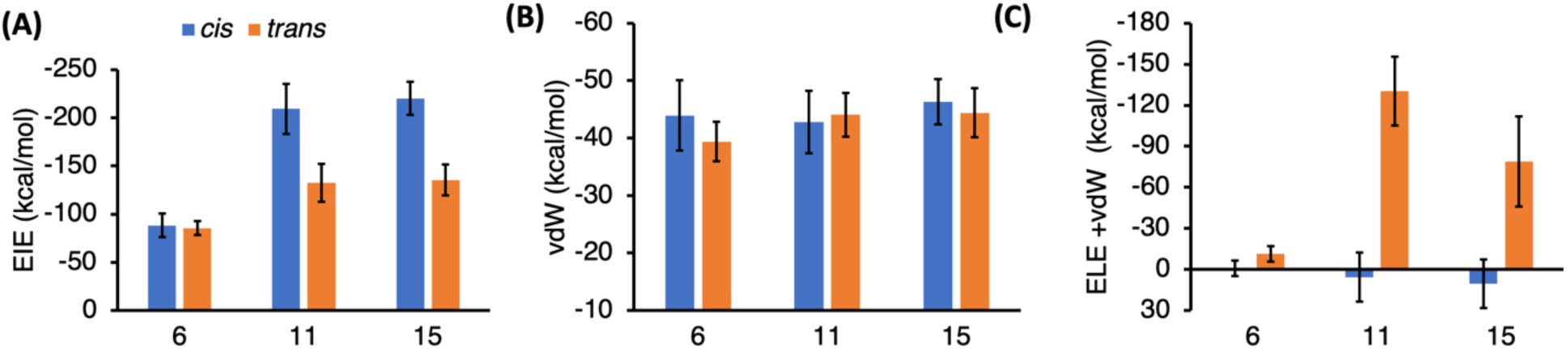
(A) Electrostatic interaction energy (EIE) and (B) van der Waals (vdW) interaction energy between eDHFR and TMP derivatives in their *cis* and *trans* isomers. (C) The sum of electrostatic and vdW ligand-water interaction energies. The ligands are in complex with eDHFR in all analyses.

**Figure 7.**
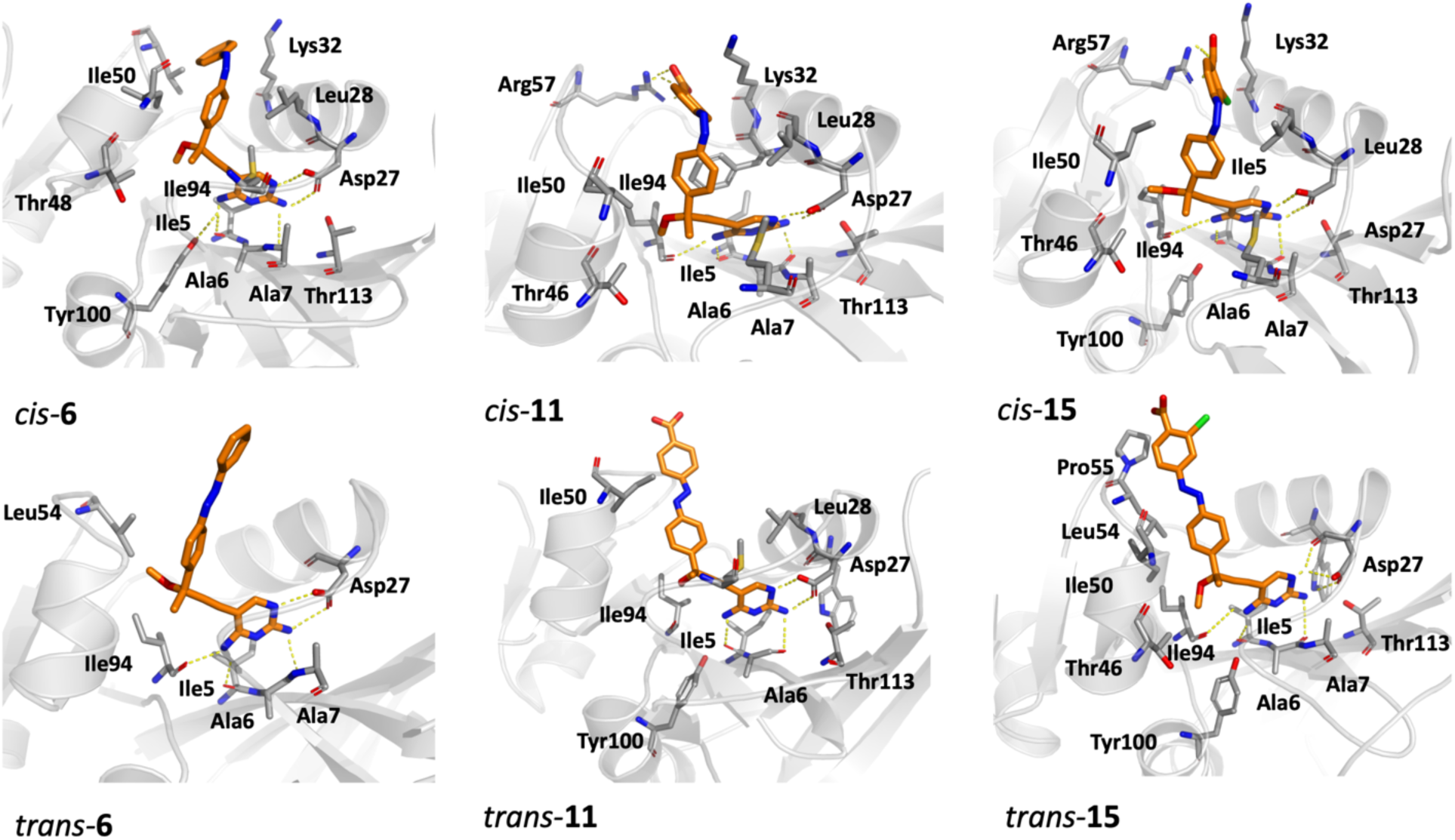
The binding poses of *cis-*and *trans*-TMP derivatives in complex with the eDHFR. The isomeric form of the ligands strongly influences their binding mode and the protein-ligand interactions in the binding site.

The energy decomposition reveals distinct interaction patterns among the three compounds. For compounds **11** and **15**, the *cis* isomers exhibit more favorable electrostatic interactions with eDHFR than the corresponding *trans* isomers, whereas compound **6** shows only a small *cis*-vs-*trans* electrostatic interaction energy difference (Figure 6A). In contrast, vdW interaction energies are relatively similar between *cis* and *trans* isomers for all three compounds (Figure 6B). These results suggest that electrostatic interactions, rather than vdW contacts, make the dominant contribution to the *cis*-vs-*trans* differential in the protein-ligand interaction energy.

For compounds **6** and **15**, the protein-ligand interaction analysis is consistent with the experimental and TI results showing “*cis*-on” effect. For compound **11**, however, the stronger electrostatic interaction of *cis*-**11** with eDHFR would appear to suggest “*cis*-on” effect, whereas both experiment and TI indicate a weak “*trans*-on” effect. This discrepancy indicates that protein-ligand interactions alone are insufficient to explain the *cis*-vs-*trans* affinity differential of compound **11**. We therefore further analyzed ligand-water interaction energies (Figure 6C). The *trans*-**11** shows the strongest stabilization from interacting with water among all systems, likely due to its greater solvent exposure in the binding pocket (Figure 7). This enhanced ligand-water interaction of *trans*-**11** can compensate for its weaker ligand-protein interactions and shift the overall binding preference toward *trans*-**11** over *cis*-**11**. By contrast, for compound **15**, the “*cis*-on” effects due to ligand-protein interaction is stronger than **11**, while “*trans*-on” effects due to ligand-water interaction is weaker than **11**. These combined effects enhance the “*cis*-on” effect of compound **15**, consistent with experiment and TI simulations.

Representative binding poses further reveals the molecular origins of these trends. For compound **6**, both isomers maintain the conserved hydrogen bond between the diaminopyrimidine N1 atom and Asp27, while *cis*-**6** forms closer contacts with Ala6, Ala7, Ile5, and Ile94. For compound **11**, Asp27 remains the primary stabilizing residue for both isomers, but the added carboxylate group strengthens electrostatic interactions in *cis*-**11** through hydrogen bond with Lys32 and Arg57. However, this protein-ligand stabilization of *cis*-**11** is offset by stronger ligand-water stabilization of trans-11. For compound 15, *cis*-**15** forms a cooperative interaction network: Asp27 stabilizes the diaminopyrimidine ring, Lys32 and Arg57 interact favorably with the carboxylate group, and Ile5, Ile94, Leu28, and Ile50 create a compact hydrophobic environment around the ligand’s aromatic rings. In *trans*-**15**, these contacts are weakened because the ligand extends away from the binding cavity. Overall, the active isomer is determined by a balance between protein-ligand electrostatics, hydrophobic packing, and ligand solvation rather than by a single residue interaction.

## Discussion

Photoswitchable ligands with titratable groups whose protonation-state equilibrium shifts in response to changes in their molecular environments represent an important class of compounds with various applications in photopharmacology. To uncover how their light-responsive binding affinities are coupled to changes in protonation, we performed a comprehensive computational investigation of TMP-derived photoswitchable inhibitors of eDHFR, developed recently through a systematic, hypothesis-driven, and structure-based experimental design campaign.^17^ The pH-REMD and *ab initio* QM/MM MD US simulations established that the ligand’s diaminopyrimidine ring is protonated upon protein binding. The protonation state change of the ligand is due to the significant increase in the pKa of this functional group, which is attributed to the strong salt bridge interaction with the Asp27 residue in the binding pocket.

Additionally, the performance of TI with different choices of alchemical transformation pathways and ligand protonation states was benchmarked against experimental data.^17^ The benchmark focused on the quantitative accuracy for predicting two critical properties for photopharmacology: (1) the *cis*-vs-*trans* affinity differential of an individual ligand, and (2) the substituents’ effects on the affinity differentials among a group of ligands. TI simulations employing a *cis*-to-*trans* alchemical transformation pathway while accounting for the protonation-state change upon protein binding (Method A) emerged as the most robust approach. Importantly, without explicitly accounting for binding-induced changes in protonation state, this approach fails to faithfully reproduce the substituents’ effects on the experimentally observed affinity differentials.

TI simulations employing compound-to-compound alchemical transformations (Method B) were less robust. When explicitly treating protonation state changes of the ligand, Method B’s performance relies on cancellations of large errors in the pKa prediction of distinct compounds if the ligand’s protonation state change is explicitly treated. The pKa errors arise from inaccuracies in the force field parameters, which typically do not cancel out between a pair of compounds connected by the alchemical transformation. When not explicitly treating the ligand’s protonation state changes, Method B cannot consistently provide qualitatively correct *cis*-vs-*trans* affinity differentials or the substituents’ effect on them.

One could, in principle, perform rigorous QM-based calculations to correct the pKa for each compound in each isomer form to remedy this issue of Method B. However, such an approach would not only incur significantly higher computational cost but may also be infeasible for large, flexible molecular photoswitches whose conformational distributions depend on the ligand’s protonation state. In our aqueous solution MD simulations, the ligand can sample many conformations, which differ in different protonation states. A simple QM-based approach employing geometry minimization and single-point energy calculation is not compatible with the large conformational ensemble of protonated and deprotonated states combined. More complex methods based on QM/MM free-energy calculations could be applied, but this would incur higher computational costs, since they need to be applied to each isomer of each ligand.

Our results also indicate that the MM-PB(GB)SA methods cannot consistently predict either the *cis*-vs-*trans* affinity differential or its substituents’ effect. Also, standard MD simulations suggest that the light-responsive affinity differential arises from a balance of protein-ligand and ligand-water interactions, whereas simply considering protein-ligand interaction can lead to qualitatively incorrect interpretations and predictions, as highlighted by the case of compound **11**. This compound exhibits a cancellation of strong “*cis*-on” and “*trans*-on” effects arising from protein-ligand and ligand-water interactions, respectively. Additionally, the light-responsive affinity differential is a result of a network of ligand-protein interactions. Persistent H-bond to key residues such as Arg57 and Lys32 stabilizes the *cis-***15** over *trans*-**15**, contributing to its strong “*cis*-on” effect.

Beyond this specific system, the methodological contribution from this work have broader implications for the design of photoswitchable ligands. The ability to computationally predict small *cis-*vs*-trans* affinity differentials and their substituents’ effect is essential for rational design of photoswitchable ligands, where energetic errors of less than 1 kcal/mol can totally reverse the light-responsive functional outcome of the biomolecular system under photocontrol. Combining multiscale free-energy methods with detailed interaction analysis, this work establishes a quantitative computational approach that links protonation state change, light-responsive binding affinity, chemical substitution, and residue-level interactions, enabling rational design principles for next-generation ionizable photoswitches in photopharmacology.

## Supplemental information

The supplemental information contains Figures S1-S2 and Table S1, as referenced throughout the main text. The figures include the MM and QM PES of the compound 6 in the vacuum, and the QM/MM partitioning scheme. The table includes the alchemical free energy changes from TI simulations. It also contains structures for the initial simulation setup of the systems.

## Author Contributions

Ruibin Liang designed the research project. Mohammad Khavani, Kambham Devendra Reddy, Pauf Neupane, Gustavo J. Costa and Laleh Khalvati performed the simulations and analyzed the data. Ruibin Liang, Mohammad Khavani, Kambham Devendra Reddy, Pauf Neupane, and Laleh Khalvati wrote and revised the manuscript.

## Acknowledgments

This work was supported by the National Institutes of Health (grant number: R35GM150780) and the Texas University Fund (TUF) for the Human Molecular Aging Center at Texas Tech University under internal award #TUF6656. We also acknowledge the computing facilities provided by the High-Performance Computing Center at Texas Tech University.

## Data and Software Availability statement

We refer the readers to the Supplemental Information file for additional data related to the properties of the molecular systems investigated in this manuscript. The structures and inputs for the initial simulation setup of the eDHFR systems are included in the “setup_files.zip” file.

