## Supplemental Information for "Multiscale Free-Energy Methods for Protonation-Coupled Light-Responsive Binding of Ionizable Photoswitchable eDHFR Inhibitors"

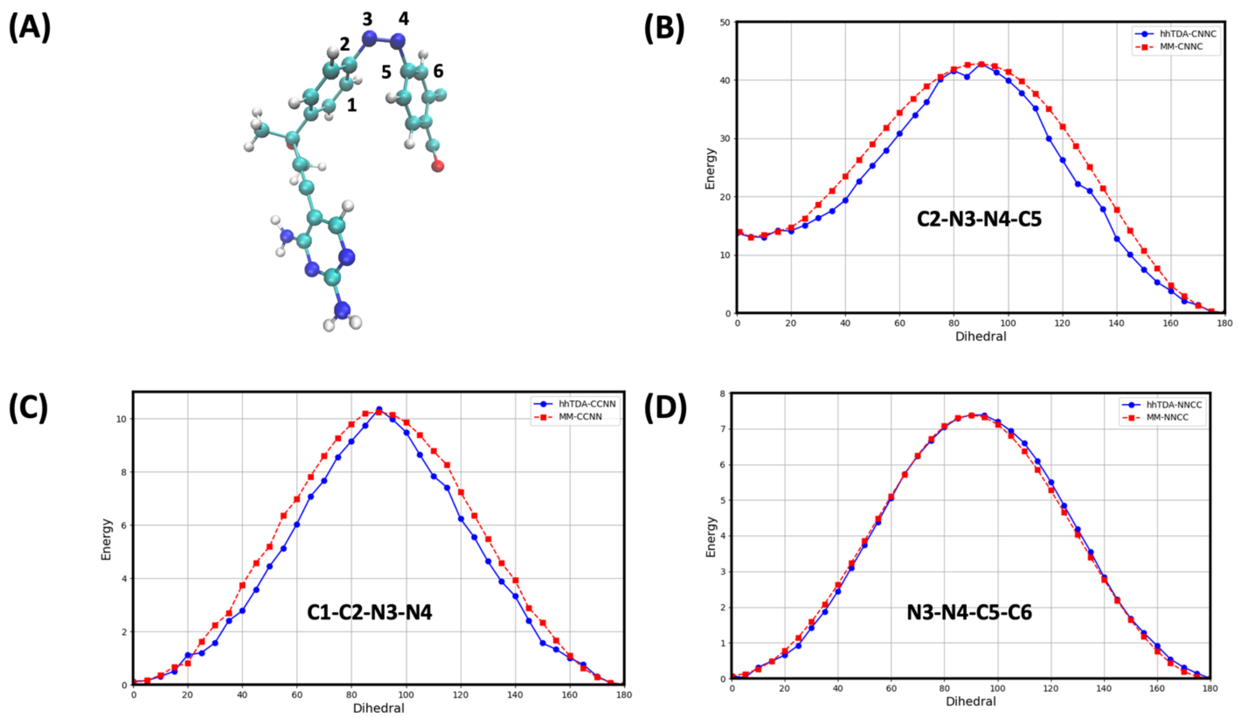


**Figure S1.** (A) Optimized ground-state geometry of the *cis*-**15** in the vacuum at the B3LYP/6-31+G(d) level of theory. (B-D) One-dimensional ground-state (PESs) for compound **15**, generated by relaxed scans along key torsions of C2-N3=N4-C5, C1-C2-N3=N4, and N3=N4-C5-C6, respectively. For each scan, the selected dihedral angle was fixed at each value, while all other degrees of freedom coordinates were fully relaxed. The multireference method, FOMO-hh-TDA-BH&HLYP/def2-SVP/6-31G+(d), was used for the QM PES scans. PES scans using the reparametrized force field (dashed red curves) are compared with the QM results (solid blue curves). For all torsions, the MM PES largely captures the shape of the QM PES, including the relative stability between the *cis* and *trans* isomers and the energy barriers separating them.

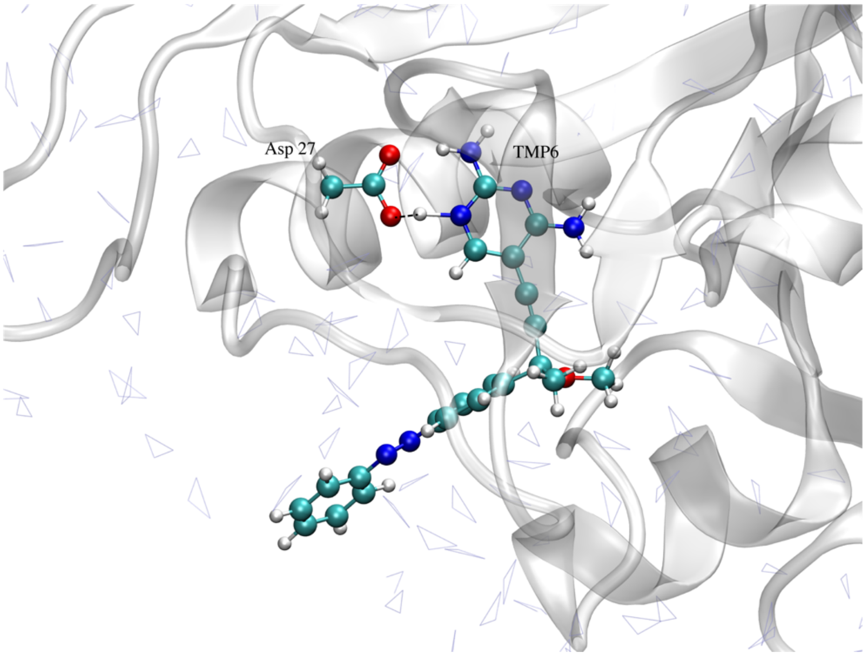


**Figure S2.** The QM region consisting of the *trans*-**6** and the side chain of the Asp27 involved in the proton transfer event. This region was treated at the B3LYP-D3/def2-SVP level of theory in the QM/MM US simulations.

**Table S1.** The *cis*-vs-*trans* affinity differentials ($\Delta\Delta G_{X,\ cis\rightarrow trans}$, in kcal/mol) and substituents' effects ($\Delta\Delta\Delta G_{X\rightarrow Y, cis-vs-trans\ binding}$ in kcal/mol) for compounds **6**, **11**, and **15**, calculated using TI (Method A). Experimental values[1] are included. Positive $\Delta\Delta G_{X,\ cis\rightarrow trans}$ values indicate "*cis*-on" effect, and vice versa. Positive $\Delta\Delta\Delta G_{X\rightarrow Y, cis-vs-trans\ binding}$ values indicate that X→Y conversion enhances the "*cis*-on" effects, and *vice versa*. The $\Delta\Delta\Delta G_{X\rightarrow Y, cis-vs-trans\ binding}$ results calculated with the same ligand protonation state in both environments are reported.

| **Quantity** | **Compounds** | | |
|---|---|---|---|
| | **6** | **11** | **15** |
| $\Delta\Delta G_{X,\ cis\rightarrow trans}$ (Exp.) | 0.5 ± 0.5 | -0.1 ± 0.4 | 0.4 ± 0.2 |
| **Deprotonated** | | | |
| $\Delta G_{unbound,\ cis\,X\rightarrow trans\,X}$ (TI) | -11.9 ± 0.1 | -11.9 ± 0.1 | -11.6 ± 0.1 |
| $\Delta G_{bound,\ cis\,X\rightarrow trans\,X}$ (TI) | -11.5 ± 0.5 | -8.7 ± 0.5 | -9.3 ± 0.9 |
| $\Delta\Delta G_{X,\ cis\rightarrow trans}$ (TI) | 0.4 ± 0.5 | 3.2 ± 0.5 | 2.3 ± 0.9 |
| **Protonated** | | | |
| $\Delta G_{unbound,\ cis\,X\rightarrow trans\,X}$ (TI) | -11.4 ± 0.2 | -10.6 ± 0.1 | -11.7 ± 0.3 |
| $\Delta G_{bound,\ cis\,X\rightarrow trans\,X}$ (TI) | -11.3 ± 0.5 | -13.0 ± 0.1 | -11.3 ± 0.1 |
| $\Delta\Delta G_{X,\ cis\rightarrow trans}$ (TI) | 0.1 ± 0.5 | -2.4 ± 0.1 | 0.4 ± 0.3 |
| | $\Delta\Delta\Delta G_{X\rightarrow Y, cis-vs-trans\ binding}$ | | |
| | **6→15** | **6→11** | **11→15** |
| **Experiment** | -0.1 ± 0.5 | -0.6 ± 0.6 | 0.5 ± 0.4 |
| **Deprotonated** | 1.9 ± 1.0 | 2.8 ± 0.7 | -0.9 ± 1.0 |
| **Protonated** | 0.3 ± 0.6 | -2.5 ± 0.5 | 2.8 ± 0.3 |
